# Metabolic regulation of ILC2 differentiation into ILC1-like cells during *Mycobacterium tuberculosis* infection

**DOI:** 10.1101/2021.01.19.427257

**Authors:** Dan Corral, Alison Charton, Maria Z Krauss, Eve Blanquart, Florence Levillain, Emma Lefrançais, Tamara Sneperger, Jean-Philippe Girard, Gérard Eberl, Yannick Poquet, Jean-Charles Guéry, Rafael J Arguello, Matthew R Hepworth, Olivier Neyrolles, Denis Hudrisier

## Abstract

Tissue-resident innate lymphoid cells (ILCs) regulate tissue homeostasis, protect against pathogens at mucosal surfaces and are key players at the interface of innate and adaptive immunity. How ILCs adapt their phenotype and function to environmental cues within tissues remains to be fully understood. Here, we show that *Mycobacterium tuberculosis* infection alters the phenotype and function of immature lung ILC2 toward a protective interferon-γ-producing ILC1-like population. This differentiation is controlled by type 1 cytokines and is associated with a glycolytic program involving the transcription factor HIF1α. Collectively, our data reveal how tissue-resident ILCs adapt to type 1 inflammation toward a pathogen tailored immune response.

## Introduction

Innate lymphoid cells (ILCs) are a population of tissue-resident cells of lymphoid origin that play a key part in both tissue homeostasis and immunity. ILCs are subdivided into three distinct populations based on their expression of cytokines and specific transcription factors. ILC1 depend on T-bet and produce interferon (IFN)-γ, ILC2 depend on GATA3 and produce interleukin (IL)-5 and IL-13, and ILC3 depend on RORγt and produce IL-17A and IL-22 (Meininger et al., 2020; Vivier et al., 2018). Based on these properties, group 1, 2 and 3 ILCs are commonly presented as the innate counterparts of T helper type 1 (Th1), Th2 and Th17 cells, contributing to type 1, 2 and 3 immune responses, respectively.

The regulome of ILCs evolves progressively during the development of each population to reach a state in which key loci specific to each lineage are acquired (Shih et al., 2016; Vivier et al., 2016). Yet, several elements controlling cytokine expression or loci encoding lineage-determining transcription factors remain broadly accessible in all ILC subsets (Shih et al., 2016). This feature contributes to the remarkable ability of ILCs to dynamically adapt to physiological or pathological alterations in their tissue of residence, and to adopt new phenotypic and functional profiles. Besides the local plasticity among mature ILC subsets (Bal et al., 2020), circulatory and tissue resident ILC precursors in human and mouse contribute to the local differentiation into mature ILCs, an *“ILCpoiesis” in situ (Lim and Di Santo, 2019)*, sustaining the ILC response depending on tissue and inflammation (Ghaedi et al., 2020; Lim et al., 2017; Zeis et al., 2020). While the various populations of tissue-resident ILCs can promptly sense and adapt to environmental changes (Meininger et al., 2020; Ricardo-Gonzalez et al., 2018) the mechanism allowing such responses remains to be fully elucidated.

In both mice and human subjects, *Mycobacterium tuberculosis* (*Mtb*) infection induces prolonged proinflammatory responses that are associated with oxidative stress, which favors tissue destruction and triggers a tissue remodeling program. *Mtb* infection is also associated with metabolic changes in the lungs, involving the utilization of aerobic glycolysis primarily instead of oxidative phosphorylation (OXPHOS) in mitochondria (Warburg effect) (Fernández-García et al., 2020; Shi et al., 2015). At the cytokine level, the lungs at steady-state mostly host resting ILC2, which, together with alveolar macrophages, imprint a type 2-oriented environment to the tissue (Saluzzo et al., 2017; Svedberg et al., 2019). *Mtb* infection of the lung triggers dramatic changes leading to the development of type 1 immunity that is mediated by IFN-γ and is associated with protection (O’Garra et al., 2013).

Here, using the murine model of *Mtb* infection, we explored how lung ILCs respond to chronic pulmonary infection and, in particular, how ILC subsets adapt and respond to type 1 inflammation within tissue. Our work uncovers the local differentiation of lung ILC2 precursors into a protective ILC1-like population through metabolic regulation during *Mtb* infection.

## Results and Discussion

### Local differentiation of ILC2 precursor into ILC1-like cells during *Mtb* infection

In order to investigate how a chronic type 1 infection impacts lung ILC subsets, C57BL/6 mice were infected with the *Mtb* reference strain H37Rv. At steady-state, *bona fide* lung ILCs were defined as a population that does not express lineage markers (CD3, CD4, CD8, TCRβ, TCRαδ, CD49b, CD11b, CD11c, B220, CD19, F4/80, GR-1, TER119, FcεR1a) but highly express CD90.2 and CD45.2 (**Figure 1A**). ILC2 (GATA3^high^), which represent the major ILC population in the murine lung, were identified after exclusion of ILC1 (NK1.1^+^) and ILC3 (RORγt^+^) cells (**Figure 1A**). At steady state, and in agreement with previously published work (Stehle et al., 2018; Vivier et al., 2018), the lung is dominantly enriched in ILC2 (**Figure 1A and 1B**). Notably, a small frequency of ILC2 expressed IL-18Rα (**Figure 1B**), a phenotype previously utilized to identify tissue-resident ILC precursor (ILCP) able to differentiate into ILC2 in the context of type 2 inflammatory responses (Ghaedi et al., 2020; Zeis et al., 2020). IL-18Rα^+^ ILC2 expressed canonical ILC2 markers such as GATA3, ST2, and Arg1 at lower levels than IL-18Rα^+^ ILC2 (**Supplementary Figure 1A-B**). At the functional level, these cells produced lower amount of IL-5 compared to IL-18Rα^−^ ILC2 and did not produce IFN-γ like ILC1 (**Supplementary Figure 1C**). In line with previous studies (Ghaedi et al., 2020; Zeis et al., 2020), we found that IL-18Rα^+^ ILC2 express high levels of TCF-1, like ILC2 precursors from the bone marrow, confirming their immature profile (**Supplementary Figure 1D**). *Mtb* infection had a profound impact on ILC composition and phenotype and was associated with gradual increase in ILC1 and ILC3 (**Supplementary Figure 1E**). Of particular interest, *Mtb* infection promoted the accumulation of a novel ILC population expressing both IL-18Rα and T-bet within the lung (**Figure 1A and 1B**) and concomitant reduction in IL-18Rα^-^ ILC2 (**Supplementary Figure 1E**). Phenotypical analysis revealed that this subset displayed little to no classical ILC2 markers, such as GATA3, ST2, Arg1 and IL-5 (**Figure 1C**), or ILC3 markers, such as RORγt (**Figure 1A**). Like ILC1, this subset expressed T-bet, CD49a, and CD226 (**Figure 1D**) but did not express NK1.1 (gated on NK1.1 negative ILCs), NKp46 or Eomes (**Figure 1A; Supplementary Figure 1E**). At the functional level, we found that this population was able to produce IFN-γ, but not IL-5 or IL-17A (**Figure 1E**). We therefore named this new subset “ILC1-like cells”, based on the similarities (**Figure 1D and 1E**) and differences (**Figure 1A; Supplementary Figure 1F**) with NK cells/ILC1. ILC1-like cells became detectable after 21 days of infection and expanded in the following weeks (**Figure 1F**). Adaptive immune responses are detectable within the lung at 21 days post-*Mtb* infection (Urdahl et al., 2011), which coincides with the detection of ILC1-like cells. As such we assessed the role of adaptive immunity in the emergence of this ILC subset. ILC1-like cells were detectable in *Mtb* infected *Rag2*^-/-^ mice that lack adaptive immunity, and at a higher level that in infected wild-type mice. Thus, adaptive immunity is not required for the generation of ILC1-like cells following infection (**Supplementary Figure 1G**).

**Figure 1.**
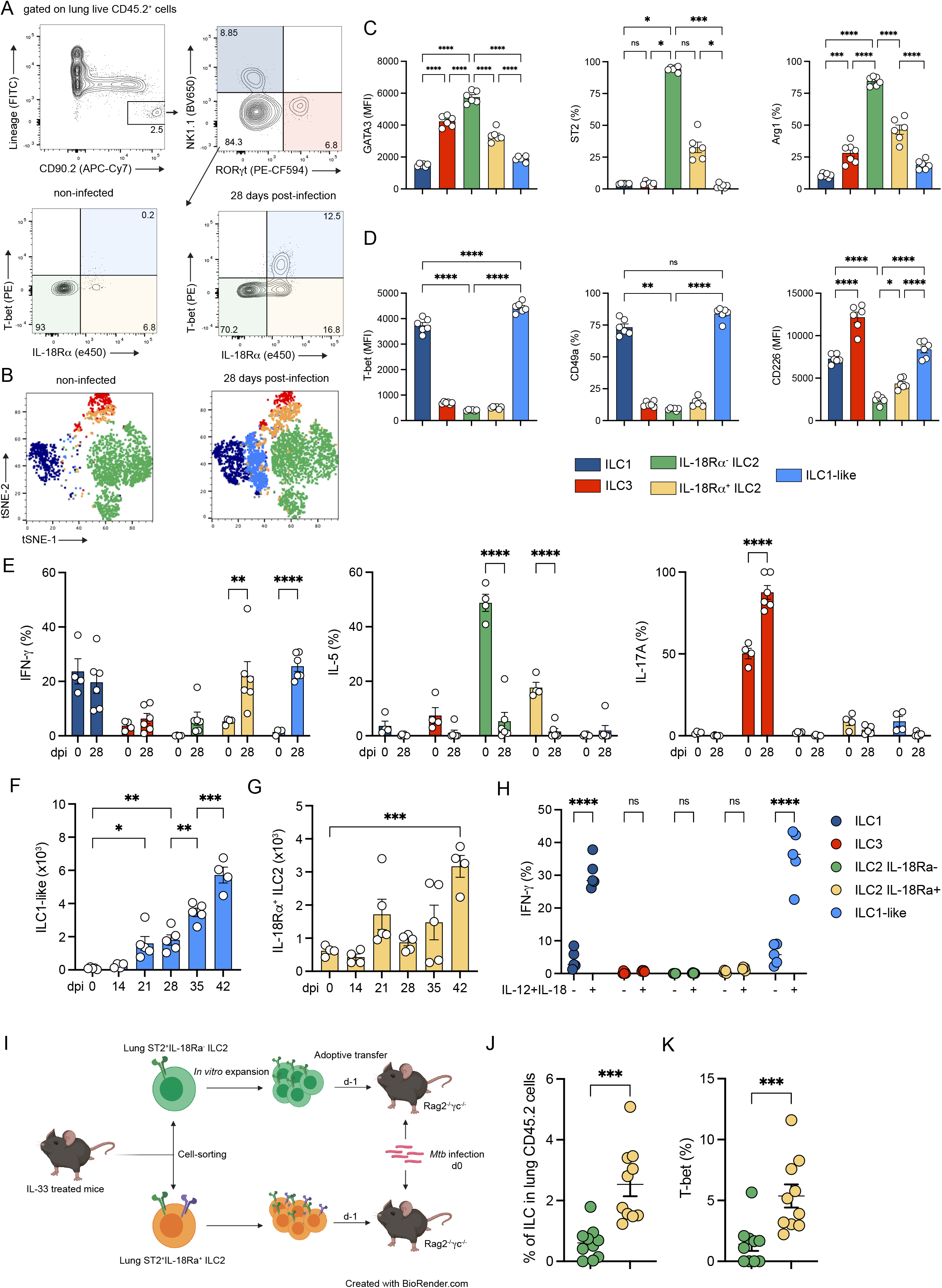
IL-18Rα–expressing ILC2 differentiate into ILC1-like cells during *Mtb* infection. **(A)** Representative dot plots showing the gating strategy used to analyze ILC subsets in the lungs of C57BL/6 mice after doublets exclusion (top graphs): ILC1 (dark blue), ILC3 (red), IL-18Rα^-^ ILC2 (green), IL-18Rα^+^ ILC2 (yellow) and T-bet^+^IL-18Rα^+^ ILC (light blue) are depicted in non-infected and *Mtb*-infected mice. **(B)** Unsupervised t-SNE distribution of total lung Lin^-^CD90.2^+^ populations at steady-state (left graph) and during *Mtb* infection (right graph). Based on gating strategy defined in Figure 1A, ILC subsets were depicted with the same color code: ILC1 in dark blue, ILC3 in red, IL-18Rα^-^ ILC2 in green, IL-18Rα^+^ ILC2 in yellow and ILC1-like cells (light blue). **(C)** Expression of GATA3 (MFI), ST2 (%), and Arg1 (%) in indicated ILC subsets at day 28 post-infection in C57BL/6 mice. **(D)** Expression of T-bet (MFI), CD49a (%) and CD226 (MFI) in indicated ILC subsets at day 28 post-infection in C57BL/6 mice. (**E**) Percentages of IFN-γ, IL-5, and IL-17A positive cells in the indicated ILC subsets after *ex vivo* stimulation with PMA/ionomycin in presence of Brefeldin A for 4h at day 28 post-infection in C57BL/6 mice. **(F)** Absolute numbers of ILC1-like cells at the indicated days after *Mtb* infection. Prior to sacrifice, mice were injected with fluorescent anti-CD45.2 to distinguish vascular and parenchymal cells. ILC1-like cells have been gated on lung-resident cells. **(G)** Percentages of IFN-γ^+^ cells in the indicated ILC subsets after *ex vivo* stimulation with IL-12+IL-18 or not, in presence of Brefeldin A for 4h at day 28 post-infection in C57BL/6 mice. **(H)** Absolute numbers of IL-18Rα^+^ ILC2 at the indicated days after *Mtb* infection. Prior to sacrifice, mice were injected with fluorescent anti-CD45.2 to distinguish vascular and parenchymal cells. IL-18Rα^+^ ILC2 have been gated on lung-resident cells. **(I)** Experimental settings for the adoptive transfer of IL-18Rα^-^ and IL-18Rα^+^ ILC2 into Rag2^-/-^γc^-/-^ before *Mtb* infection. **(J)** Percentage of ILC (Lin^-^CD45.2^+^CD90.2^+^CD127^+^) in lung at day 21 post-infection in Rag2^-/-^γc^-/-^ after adoptive transfer of IL-18Rα^-^ ILC2 (green) and IL-18Rα^+^ ILC2 (yellow). **(K)** As in **(J)**, but for T-bet expression in ILC. In **(C, D, F and H)**, data are representative of five independent experiments, in **(E, G)** data are representative of two independent experiments and in **(J, K)** data are a pool of two independent experiments with each symbol representing an individual mouse, graphs depict data as mean (± s.e.m) and statistical analysis was performed using two-way **(E-F)**, one-way ANOVA **(C-D, G-H) or** Mann Whitney test **(J-K)** (*, p<0.05; **, P<0.01; ***, p<0.001; ****, p<0.0001).

ILCs have been reported to adapt their profile to environmental cues. Different mechanisms have been described to sustain the local adaptation of ILCs during inflammation such as plasticity of mature ILC subsets (Bal et al., 2020), *in situ* differentiation of ILC precursor (Ghaedi et al., 2020; Zeis et al., 2020) as well as the migration of ILC from bone marrow (Zeis et al., 2020). Furthermore, ILCs with characteristics of ILC1-like cells have been shown to arise from various ILC subsets through mechanisms of plasticity depending on the tissue and the inflammatory context (Bal et al., 2020; Silver et al., 2016). In the lungs, ILC2 have been described to acquire expression of T-bet, IL-18Rα and IFN-γ during influenza virus infection (Silver et al., 2016). We hypothesized that ILC1-like cells could differentiate from lung ILC2. To assess if ILC2 display plasticity during *Mtb* infection, we adoptively transferred total lung ST2^+^ ILC2, sorted regardless of their IL-18Rα expression, into *Rag2*^-/-^*Il2rg*^-/-^ mice, which are devoid of T cells, B cells and NK/ILCs, one day prior to *Mtb* infection (**Supplementary Figure 1H and 1I**). Before transfer, we confirmed that sorted ILC2 expressed GATA3 but not T-bet or RORγt (**Supplementary Figure 1J**) and noticed that IL-18Rα expression was lost during *in vitro* culture (**Supplementary Figure 1J**). Following transfer, ILC2 strongly upregulated T-bet in infected, but not in non-infected mice (**Supplementary Figure 1K and 1L**). Furthermore, T-bet^high^ cells expressed higher level of IL-18Rα and Ki67 compared to GATA3^high^ cells (**Supplementary Figure 1M**). Given that ILC2 can give rise to ILC1-like cells, we sought to explore which ILC2 subset preferentially differentiated into ILC1-like cells. Intriguingly, while IL-18Rα^+^ ILC2 did not acquire ILC1 markers during *Mtb* infection (**Figure 1D**) and accumulate into the lungs (**Figure 1G**), they gained the potential to produce IFN-γ and did not produce IL-5 (**Figure 1E**). Moreover, IL-18Rα^+^ ILC2, unlike ILC1 and ILC1-like cells, did not respond to *ex vivo* stimulation with IL-12 and IL-18 (**Figure 1H**). Thus, we hypothesized that this population could have the potential to differentiate into ILC1-like cells and could thus represent an early stage of ILC1-like. We assessed if IL-18Rα^+^ ILC2 have the potential to differentiate into ILC1-like during *Mtb* infection. To this end, we sorted ILC2 subsets from IL-33 treated mice based on their IL-18Rα expression and adoptively transferred them into *Rag2*^-/-^*Il2rg*^-/-^ mice one day before infection with *Mtb* (**Figure 1I**). Interestingly, we found that the transfer of IL-18Rα^+^ ILC2 resulted in higher proportions of ILC in the lungs when compared to the conditions where the same number of IL-18Rα^-^ ILC2 were transferred (**Figure 1J**); in addition, T-bet expression among IL-18Rα^+^ ILC2 was significantly increased in the former case (**Figure 1K**). Thus, IL-18Rα^+^ ILC2, rather than IL-18Rα^-^ ILC2, have the potential to differentiate into ILC1-like cells during *Mtb* infection.

Altogether, our data show that *Mtb* infection differentially impacts the composition of ILC subsets within the lung, and especially induces the local differentiation of lung ILC2 precursor into a ILC1-like cell population.

### Type 1 inflammatory environment shapes the fate of IL-18Rα^+^ ILC2

Next, we aimed to assess how the inflammatory milieu influences the fate of IL-18Rα^+^ ILC2. *Mtb* infection triggers the development of a type 1 immunity (O’Garra et al., 2013). Both IL-12 and IL-18 contribute in the establishment of this type 1 inflammatory environment (Kinjo et al., 2002; O’Garra et al., 2013) in particular by inducing the expression of IFN-γ on ILC1, NK cells and Th1 (Chiossone et al., 2018; Weizman et al., 2017). Therefore, we administered to mice IL-12 and IL-18 intranasally for 1 week and found that this treatment was sufficient to induce the accumulation of ILC1-like cells in the lungs (**Figure 2A-B**). Furthermore IL-18Rα^+^ ILC2 also expanded in these settings (**Figure 2B**) and lost the expression of TCF-1 (**Figure 2C**), supporting the idea that these cells may underwent a local differentiation process. Similarly, to *Mtb* infection, IL-18Rα^+^ ILC2, and IL-18Rα^-^ ILC2, lost their ability to produce IL-5 following cytokine injection and acquired the ability to produce IFN-γ (**Figure 2D**), but not after *ex vivo* stimulation with IL-12 and IL-18 (**Figure 2E**). Moreover, we crossed IL-5^Cre-tdTomato^ (Red5) mice with ROSA26-YFP mice to enable fate-mapping mature ILC2 (Nussbaum et al., 2013) and that the majority of ILC1-like cell do not derive from IL-18Rα-ILC2, the only ILC population expressing IL-5 at steady-state and during type 1 inflammation (**Supplementary Figure 2A-B**), reinforcing our previous observation (**Figure 1J**). Overall, based on the expression of several markers (GATA3, Arg1, T-bet, IL-18Rα, CD49a, CD226 and Ki67), we found close similarities in both IL-18Rα^+^ ILC2 and ILC1-like cells generated upon either IL-12/IL-18 treatment or during *Mtb* infection (**Supplementary Figure 2B**). Thus, the generation of ILC1-like cells observed during *Mtb* infection can be closely recapitulated with the simple administration of IL-12 and IL-18.

**Figure 2.**
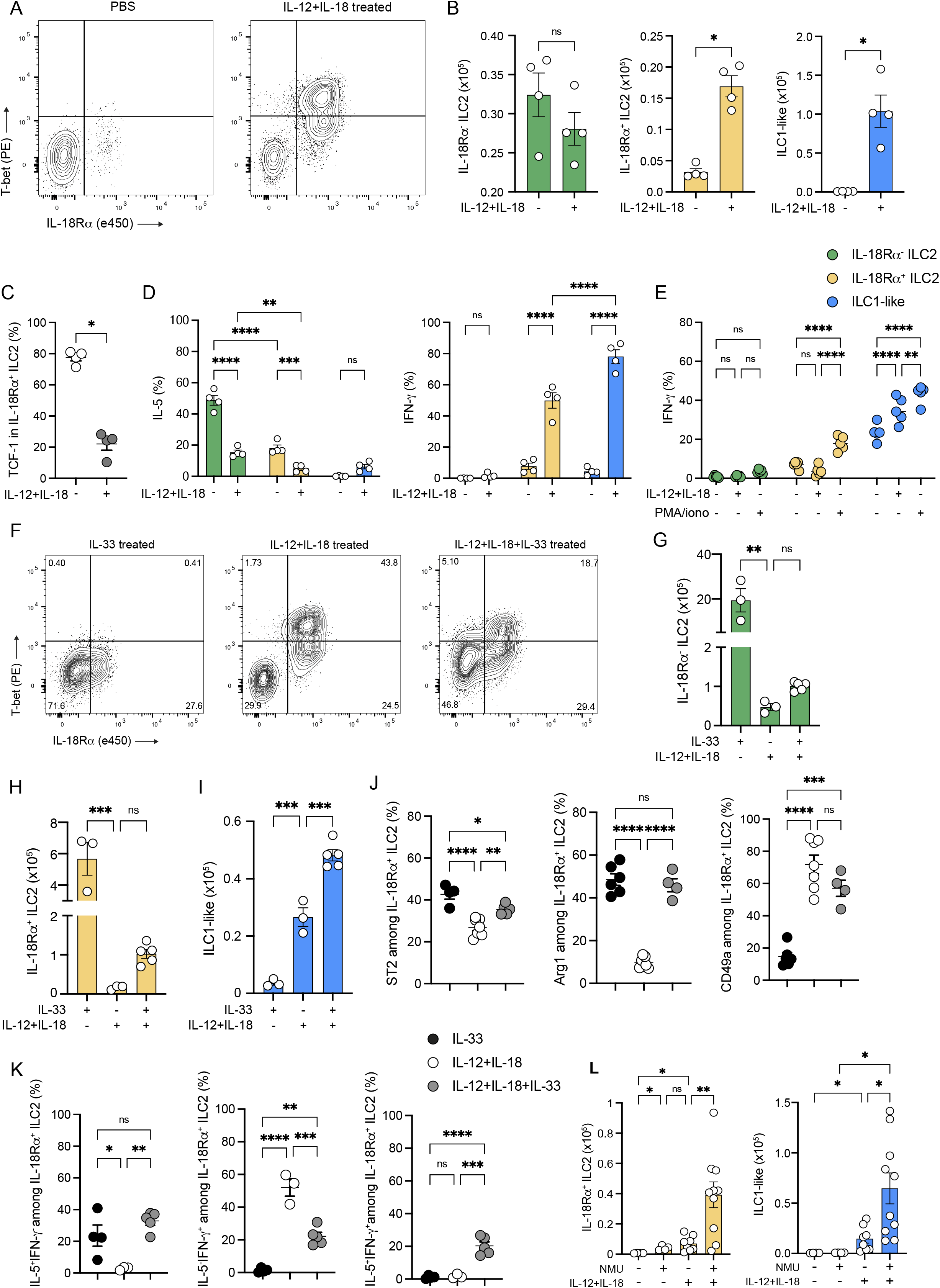
The inflammatory environment shapes the fate of IL-18Rα^+^ ILC2. **(A)** Representative dot-plots of T-bet and IL18Rα expression after intranasal administration of PBS and IL-12+IL-18 in *Rag2*^-/-^ mice in Lin^-^CD45.2^+^CD90.2^+^NK1.1^-^ RORγt^-^ cells **(B)** Absolute numbers of IL-18Rα^-^ ILC2 (green), IL-18Rα^+^ ILC2 (yellow) and ILC1-like cells (blue) after cytokine (IL-12+IL-18) or control (PBS) treatment. **(C)** Percentage of TCF-1 in lung IL-18Rα^-^ ILC2 after intranasal administration of PBS (white dots) or IL-12+IL-18 (grey dots). **(D)** Percentages of cells expressing IL-5 or IFN-γ among the indicated ILC subsets after *ex vivo* stimulation with PMA/ionomycin in the presence of brefeldin A for 4h. **(E)** Percentages of IFN-γ^+^ cells in the indicated ILC subsets after *ex vivo* stimulation with IL-12+IL-18, PMA/ionomycin or not in presence of Brefeldin A for 4h from IL-12+IL-18 treated C57BL/6 mice. **(F)** Representative dot-plots of T-bet and IL18Rα expression after intranasal administration of IL-33, IL-12+IL- 18 and IL-12+IL-18+IL-33 in *Rag2*^-/-^ mice in Lin^-^CD45.2^+^CD90.2^+^NK1.1^-^RORγt^-^ cells. **(G-I)** Absolute numbers of IL-18Rα^-^ ILC2 **(G)**, IL-18Rα^+^ ILC2 **(H)** and ILC1-like cells **(I)** after intranasal administration of IL-33, IL-12+IL-18, or IL-12+IL-18+IL-33. **(J)** Expression of ST2 (%), Arg1 (%) and CD49a (%) in IL-18Rα^+^ ILC2 in IL-33 (black dots), IL-12+IL-18 (white dots) and IL-12+IL-18+IL-33 (grey dots) -treated mice. **(K)** Percentage of IL-5^+^IFN- γ^-^ (right), IL-5^-^IFN- γ^+^ (middle), IL-5^+^IFN- γ^+^ (left) in IL-18Rα^+^ ILC2 in IL-33 (black dots), IL-12+IL-18 (white dots) and IL-12+IL-18+IL-33 (grey dots) -treated mice. **(L)** Absolute numbers of IL-18Rα^+^ ILC2 (left) and ILC1-like cells (right) after intranasal administration of PBS (control), neuromedin U (NMU), IL-12+IL-18, or IL-12+IL-18+NMU. Each symbol represents an individual mouse. Statistical analysis was performed using Mann-Whitney **(B, C)**, one-way **(G-K)** and two-way **(D-E, L)** ANOVA tests (*, p<0.05; **, P<0.01; ***, p<0.001; ****, p<0.0001). Graphs depict data as mean (± s.e.m). Data are representative of three **(B, D, G-K)**, and two **(C, E, L)** independent experiments.

To further demonstrate the ILC2 origin of ILC1-like cells during type 1 inflammation, we studied the effect of IL-33, a well-known inducer of both mature and immature ILC2 (Moro et al., 2010; Neill et al., 2010; Price et al., 2010), on ILC1-like differentiation. IL-33 alone did not induce the differentiation of ILC1-like cells, although it did induce a high expansion of mature and immature ILC2 (**Figure 2F-I**). In association with IL-12 and IL-18, IL-33 was able to enhance ILC1-like differentiation (**Figure 2I** and **Supplementary Figure 2C)**. Intriguingly, while IL-18Rα^+^ ILC2 expressed ILC2 markers (ST2, Arg1 and IL-5) in IL-33-treated mice (**Figure 2J-K**), and ILC1-like markers (CD49a, IFN-γ) in IL-12/IL-18-treated animals (**Figure 2J-K**), the combination of IL-12/IL-18 with IL-33 led to a mixed ILC1/ILC2 phenotype with the capacity to produce both IL-5 and IFN-γ. Because ST2 is expressed by various cell types, including ILC2, we also tested Neuromedin U (NMU), whose receptor is solely present in bone marrow ILC2P and in lung ILC2 (Cardoso et al., 2017; Klose et al., 2014; Wallrapp et al., 2017). Similar to IL-33, NMU potentiated the differentiation of ILC1-like cells induced by IL-12 and IL-18 (**Figure 2L**). Altogether, these results demonstrate that lung IL-18Rα^+^ ILC2 exhibit a highly adaptable phenotype, dependent on the inflammatory environment. While they strengthen the ILC2 response in a type 2 environment (*i.e.*, after administration of IL-33), these cells rather differentiate into IFN-γ-producing ILC1-like cells in a type 1 environment (*i.e.*, after administration of IL-12 and IL-18). This result supports the recent notion that local ILC precursors may undergo “*ILCpoiesis*” (Ghaedi et al., 2020; Zeis et al., 2020), as demonstrated in human (Lim et al., 2017; Lim and Di Santo, 2019). Although we cannot exclude local plasticity of other ILC subsets, or ILCP recruitment from the bone marrow, our results strongly suggest the local differentiation of lung ILC2P into ILC1-like cells during type 1 inflammation.

### ILC1-like cell differentiation is associated with a metabolic reprogramming toward glycolysis

Recent RNA-sequencing analyses of intestinal ILCs revealed that each subset display unique metabolic profiles (Gury-BenAri et al., 2016). While the need in amino acid metabolism for lung ILC2 functions relies on Arg1 (Monticelli et al., 2016), the glycolytic pathway necessary for ILC3 functions depends on mTOR and HIF1α (Di Luccia et al., 2019). However, little is known about metabolic adaptation of ILCs to their environment during infection (Joseph et al., 2018). Fate decisions of immune cells such as those underlying differentiation of Treg/Th17 or Treg/Th1 have been tightly associated with metabolic reprogramming (Clever et al., 2016; Dang et al., 2011; Shi et al., 2011). Given that IL-18Rα^+^ ILC2 present the ability to differentiate into ILC1-like cells in a type 1 inflammatory context, we investigated the metabolic pathways engaged during ILC1-like cells differentiation. To gain insight in ILC metabolism, we took advantage of the recently described SCENITH method (Argüello et al., 2020), which allows to determine global metabolic dependencies and capacities at the single cell level. SCENITH uses protein synthesis levels as a readout and is particularly appropriate to analyze the metabolism of rare cells, such as ILCs. ILC1-like cells were compared to control cells known to rely on a glycolytic metabolism (*e.g.,* NK cells) and to ILC2 in lungs. In agreement with the inhibitory effect of type 1 inflammation on IL-18Rα^-^ ILC2 (**Figure 1E; Supplementary Figure 1F**), administration of IL-12 and IL-18 downregulated ILC2 global level of translation, as assessed via detection of puromycin incorporation (**Figure 3A-C**). Conversely, following cytokine injection, the level of translation was increased in NK cells, IL-18Rα^+^ ILC2 and ILC1-like cells with the latter cells displaying the highest rate (**Figure 3A-C**). Notably, a similar pattern was observed during *Mtb* infection (**Supplementary Figure 3A**). The analysis of protein synthesis in the presence of inhibitors targeting different metabolic pathways, namely 2-DG for glycolysis and oligomycin for OXPHOS, allowed us to assess the mitochondrial dependence and glycolytic capacity of the cells (**Figure 3D**). We found that, in all ILC subsets tested, type 1 inflammation led to a global decrease in their mitochondrial dependence, together with an increase in their glycolytic capacity, that is a canonical feature of the Warburg effect (Heiden et al., 2009) (**Figure 3E and F**). Thus, while a metabolic reprogramming towards glycolysis is significantly induced in IL-18Rα^-^ ILC2, IL-18Rα^+^ ILC2 and characterized ILC1-like cells upon type 1 inflammation, this program was also associated with a global inhibition of IL-18Rα^-^ ILC2 compared to the other ILC subsets tested. Arg1 has been previously identified as a critical component of the metabolic programming of lung ILC2, with its inhibition or genetic inactivation resulting in reduced aerobic glycolysis (Monticelli et al., 2016). In agreement with previous studies (Bando et al., 2013; Monticelli et al., 2016; Schneider et al., 2019), we found that Arg1 was highly expressed in both IL-18Rα^-^ and IL-18Rα^+^ ILC2 (**Figure 3G**). However, after treatment with IL-12 and IL-18, the expression of Arg1 decreased in IL-18Rα^+^ ILC2 to similar level as ILC1-like cells, supporting the idea that Arg1 is not implicated in the metabolic regulation of ILC1-like cell differentiation. Previous work proposed that the Warburg effect, a metabolic pathway that is engaged during *Mtb* infection (Fernández-García et al., 2020; Shi et al., 2015), relies on the transcription factor hypoxia-inducible factor-1 α (HIF1α) (Palazon et al., 2014). Of interest, in a model of von Hippel-Lindau (VHL) deficiency, where HIF1α is overexpressed, it was previously shown that ILC2 development was repressed through glycolysis induction (Li et al., 2018). These observations prompted us to analyze HIF1α expression in our model. Type 1 inflammation driven by IL-12/IL-18 treatment led to the induction of HIF1α in both IL-18Rα^-^ and IL-18Rα^+^ ILC2, and in ILC1-like cells (**Figure 3H**). Given that IL-18Rα^-^ and IL-18Rα^+^ ILC2 share the same metabolic dependence but differ in their activation state during type 1 inflammation, we analyzed the impact of HIF1α expression on ILC2. We performed an *in vitro* assay using sorted ILC2 cultured in the presence of DMOG, which stabilizes the HIF1α protein (Palazon et al., 2014) (**Figure 3I**). DMOG-treatment had a significant inhibitory impact on expression of ILC2 markers including GATA3, ST2 and IL-5 (**Figure 3J-L**). Accordingly, analysis of the global metabolic profile of ILC2 revealed that DMOG-treated ILC2 harbored a glycolytic profile (**Figure 3M**), while untreated ILC2 preferentially used mitochondrial respiration (**Figure 3N**). Most notably, DMOG treatment alone was sufficient to upregulate genes typically associated with an ILC1 phenotype, such as *Tbx21*, *Ifng* and *Il18r1* (**Figure 3O**). Altogether, these results suggest that HIF1α in lung ILC2 promote a metabolic reprogramming toward glycolysis, while favoring the acquisition of an ILC1-like profile.

**Figure 3.**
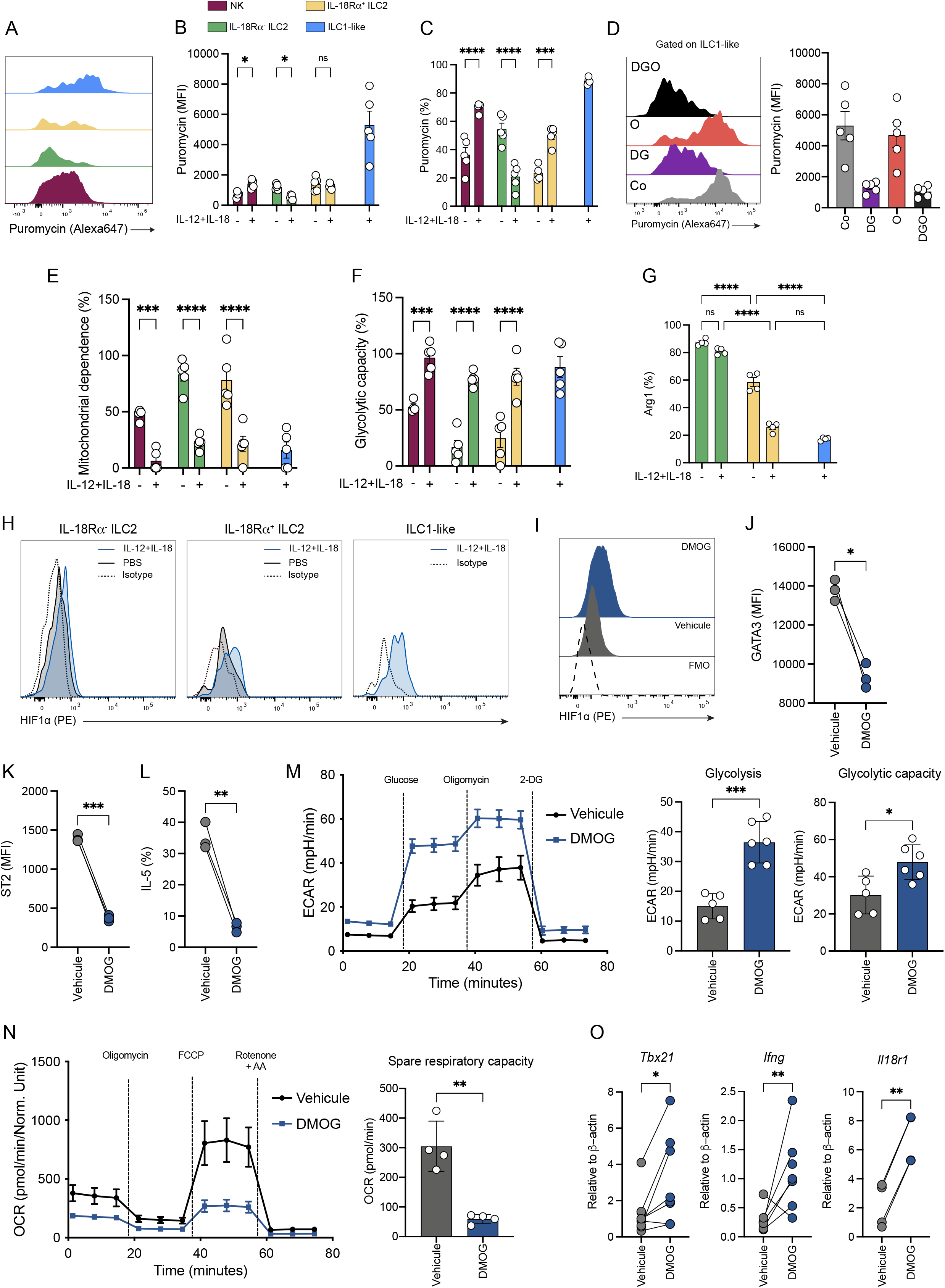
Metabolic reprogramming toward glycolysis is associated with an ILC1-like cell differentiation. **(A)** Representative histograms of puromycin staining in NK cells (violet), IL-18Rα^-^ ILC2 (green), IL-18Rα^+^ ILC2 (yellow), and ILC1-like cells (blue) in IL-12+IL-18-treated mice. **(B)** Expression of puromycin (MFI) in NK cells (violet), IL-18Rα^-^ ILC2 (green), IL-18Rα^+^ ILC2 (yellow), and ILC1-like cells (blue) in PBS *vs.* IL-12+IL-18 treated mice. **(C)** Percentages of puromycin-positive cells in NK cells (violet), IL-18Rα^-^ ILC2 (green), IL-18Rα^+^ ILC2 (yellow), and ILC1-like cells (blue) in PBS *vs.* IL-12+IL-18 treated mice. **(D)** Representative histograms of puromycin staining (left) and quantification (MFI, right) in ILC1-like cells from IL-12+IL-18 treated mice after incubation with various metabolic inhibitors (Co, control; DG, 2-Deoxyglycose; O, oligomycin; DGO, 2-Deoxyglucose + Oligomycin). **(E-F)** Percentages of mitochondrial dependence **(E)**, and glycolytic capacity **(F)** in NK (violet), IL-18Rα^-^ ILC2 (green), IL-18Rα^+^ ILC2 (yellow), and ILC1-like cells (blue) in PBS *vs.* IL-12+IL-18 treated mice. **(G)** Percentages of Arg1^+^ cells among IL-18Rα^-^ ILC2 (green), IL-18Rα^+^ ILC2 (yellow), and ILC1-like cells (blue) defined after intranasal administration of PBS (control) or cytokines (IL-12+IL-18). **(H)** Histograms showing HIF1α expression in IL-18Rα^-^ ILC2 (left), IL-18Rα^+^ ILC2 (middle) and ILC1-like cells in PBS (black line) or IL-12+IL-18 (blue line) treated mice. Dash line represents isotype control for HIF1α staining. Due to the absence of ILC1-like cells at steady-state, only the condition with IL-12+IL-18 treatment is represented. **(I)** Histograms showing HIF1α protein expression in ILC2 cultured in the absence (vehicle, grey) or presence of DMOG (blue). The dot line represents FMO for HIF1α detection. **(J-L)** Quantitative analysis of the intensity of GATA3 **(J)**, ST2 **(K)** and IL-5 **(L)** expression in ILC2 cultured in the absence (vehicle) or presence of DMOG. **(M)** Seahorse analysis of glycolytic stress test with quantification of glycolysis and glycolytic capacity of ILC2 cultured in the presence or absence of DMOG. **(N)** Seahorse analysis of mitochondrial respiration with quantification of spare respiratory capacity of ILC2 cultured in the presence or absence of DMOG. **(O)** as in **(J-L)** except that the expression of *Tbx21*, *Ifng* and *Il18ra1* mRNA was analyzed by RT-qPCR. Each symbol represents an individual mouse and statistical analysis was performed using two-way ANOVA **(B, C, E, F, G)**, and paired t test **(J-O)** (*, p<0.05; **, P<0.01; ***, p<0.001; ****, p<0.0001). Graphs depict data as mean (± s.e.m) from two **(A-N)** and a pool of three **(O)** independent experiments.

Next, we sought to determine the role of glycolysis in the differentiation of ILC1-like cells during *Mtb* infection. First, we observed that HIF1α is expressed in all ILC subsets, with the highest levels in ILC3 and ILC1-like cells (**Supplementary Figure 3B**). Inhibition of glycolysis during *ex vivo* stimulation of total lung ILCs decreased the proportion of IFN-γ^+^ ILCs (**Supplementary Figure 3C**), showing that IFN-γ production is glycolysis-dependent. To assess the impact of glycolysis induction *in vivo*, we first treated mice with 2-deoxyglucose (2-DG), a glycolysis inhibitor, during *Mtb* infection. 2-DG administration markedly decreased the number of ILC1-like cells as well as their ability to produce IFN-γ (**Supplementary Figure 3D, E**). Next, since glucose is consumed in the lungs of *Mtb*-infected mice (Fernández-García et al., 2020), we investigated if the modulation of glucose availability in the lung environment could modulate the differentiation of ILC1-like cells. Glucose supplementation in the animals’ drinking water enhanced the differentiation of ILC1-like cells (**Supplementary Figure 3F**) and increased the percentage of IFN-γ^+^ ILC1-like cells (**Supplementary Figure 3G**). Thus, these results strongly suggest that glycolysis is required to support ILC1-like cells differentiation and function during *Mtb* infection.

### ILC1-like cells confer protection against *Mtb*

Next, we investigated whether BCG, the only available vaccine for TB, might impact the population of lung-resident ILCs when delivered intranasally, a route providing a better protection than the conventional subcutaneous route (Perdomo et al., 2016), prior to *Mtb* infection (**Figure 4A**). As expected, mucosal BCG vaccination induced protection upon *Mtb* challenge (**Figure 4B**). In vaccinated mice, protection correlated with an increase in T-bet expression in ILCs (**Figure 4C**). More importantly, although BCG vaccination had no impact on other ILC subsets (**Figure 4D**), higher numbers of ILC1-like cells were detected at 14 days post-infection, a time when IFN-γ-producing ILC1-like cells were virtually absent from non-vaccinated mice (**Figure 4D, E**) but well-induced in vaccinated animals (**Figure 4D, E**). Overall, BCG vaccination promotes ILC1-like cells in early stages of infection, which could contribute to protection against *Mtb*.

**Figure 4.**
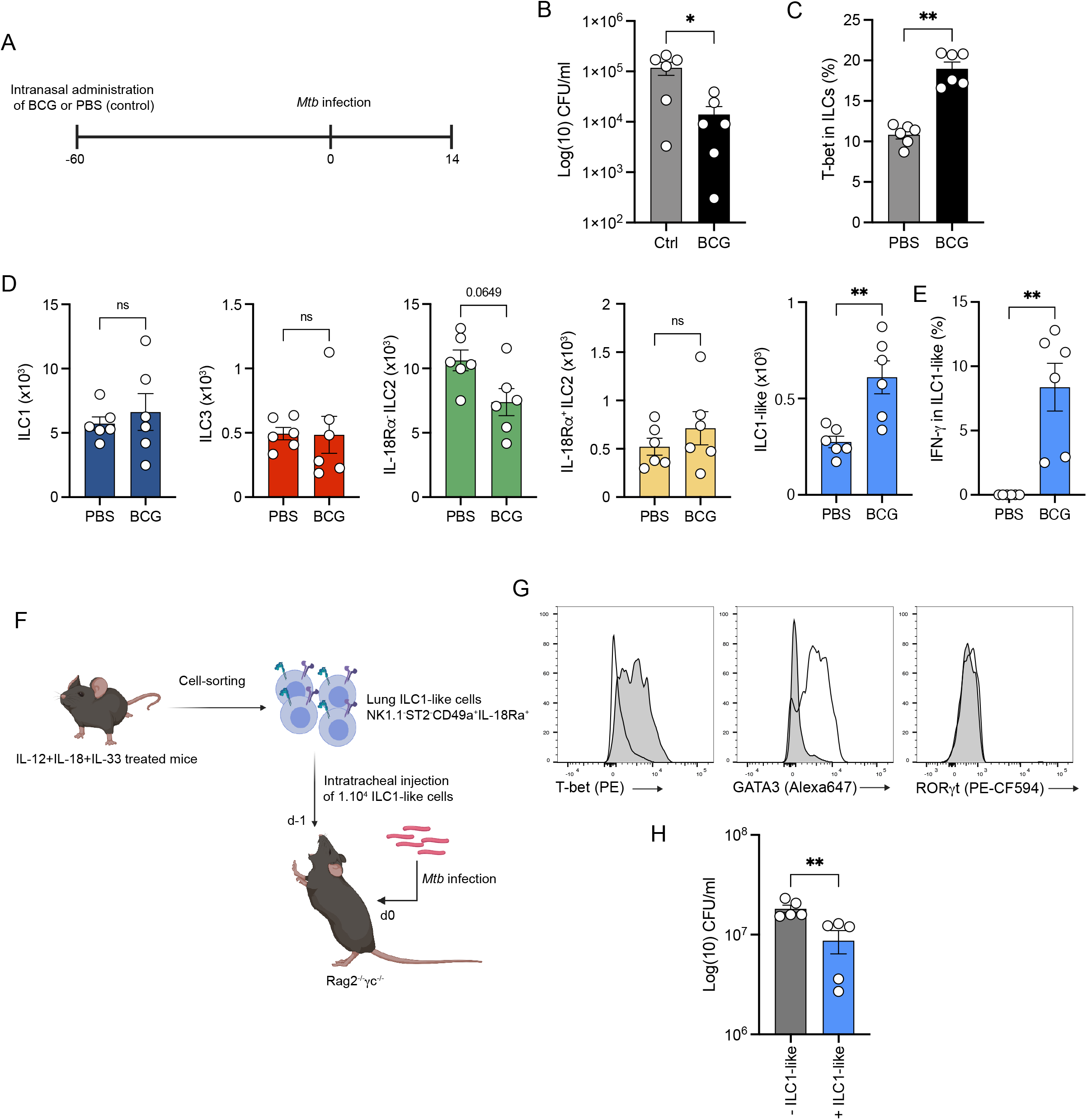
ILC1-like cells confer protection against *Mtb*. **(A)** C57BL/6 were vaccinated by internasal administration of BCG or not (PBS) 60 days before infection. After 14 days post-infection, mice were euthanized. **(B)** Mycobacterial loads at day 14 post-infection in C57BL/6 mice vaccinated or not (PBS) with 1×10^5^ BCG via the intranasal route 60 days prior *Mtb* infection. **(C)** Percentages of total lung ILC expressing T-bet. **(D)** Absolute numbers of ILC1 (dark blue), ILC3 (red), IL-18Rα^-^ ILC2 (green), IL-18Rα^+^ ILC2 (yellow) and ILC1-like cells (light blue) at day 14 post-infection in C57BL/6 mice vaccinated or not (PBS) with 1×10^5^ BCG via the intranasal route 60 days prior *Mtb* infection. **(E)** Percentages of IFN-γ^+^ cells among ILC1-like cells in *Mtb*- infected unvaccinated *vs.* vaccinated mice after *ex vivo* stimulation with IL-12+IL-18 in the presence of brefeldin A for 4h. **(F)** Schematic representation of the *in vivo* expansion of ILC1-like cells in *Rag2*^-/-^ mice treated with IL-12+IL-18+IL-33, cell-sorting of ILC1-like cells (Lin^-^CD45.2^+^CD90.2^+^NK1.1^-^ST2^-^CD49a^+^IL-18Rα^+^) and adoptive transfer in *Rag2*^-/-^*Il2rg*^-/-^ one day before infection with *Mtb* by intratracheal route. **(G)** Representative histograms of T-bet, GATA3 and RORγt expression in sorted ILC1-like cells (grey) *vs.* ILC2 (Lin^-^CD45.2^+^CD90.2^+^NK1.1^-^ST2^+^ cells). **(H)** Bacterial loads at day 21 post-infection in *Rag2*^-/-^*Il2rg*^-/-^ mice having received (+ILC1-like) or not (-ILC1like) an adoptive transfer of ILC1-like cells from IL-12+IL-18+IL-33 treated *Rag2*^-/-^ mice one day before *Mtb* infection. Each symbol represents an individual mouse. Statistical analysis was performed using Mann-Whitney test **(B-H)** (*, p<0.05; **, P<0.01; ***, p<0.001; ****, p<0.0001). Graphs depict data as mean (± s.e.m). Data are representative of two **(B-H)** independent experiments.

Finally, to assess the contribution of ILC1-like cells to protection against *Mtb*, we took advantage of the cytokine-induced ILC1-like cell model (**Figure 4F**) to generate enough ILC1-like cells for adoptive transfer. ILC1-like cells were sorted from the lungs of mice treated with IL-12, IL-18 and IL-33; these cells expressed T-bet, but not GATA3 or RORγt (**Figure 4G**). Remarkably, the transfer of as few as 10,000 ILC1-like cells resulted in a statistically significant reduction in bacterial load after *Mtb* challenge, demonstrating the protective capacity of ILC1-like cells against the tuberculosis bacillus (**Figure 4H**). Recently, ILC3 were reported to mediate protection against *Mtb* through induction of lung ectopic lymphoid follicles (Ardain et al., 2019). Although our results confirm the expansion and activation of ILC3 during *Mtb* infection (**Figure 1E** **; Supplementary Figure 1F)**, we report the expansion of an ILC1-like cell population, which can contribute to protection against *Mtb* infection. Differences between the two studies may be due to the *Mtb* strains used, namely HN878 (Ardain et al., 2019) *vs.* H37Rv in our study, the different proportions of the various ILC subsets found in the lungs, or both. In particular, infection with the hypervirulent strain HN878 is known to induce a different inflammatory pattern (*e.g.* with strong production of IL-1β and type 1 IFN) and protective immune mechanisms (*e.g.* IL-17 and IL-22 production) compared to H37Rv (Gopal et al., 2014; Manca et al., 2001). Thus, depending on the strain and the associated inflammation, ILC subsets might be highly impacted in their regulation and function during infection.

Thus, we propose a model in which the local differentiation of lung ILC2 precursor into ILC1-like cells is regulated by both inflammatory and metabolic environment induced by *Mtb* infection (**Supplementary Figure 4).** Our observation that BCG vaccination favors the early generation of ILC1-like cells and that ILC1-like cell are endowed with a protective potential during *Mtb* infection lead to future studies aiming at elucidating the role played by ILC1-like cells in protection. On a broader perspective, targeting ILC1-like cells using dedicated strategies may help develop novel approaches to combat tuberculosis and other inflammatory diseases.

## Materials & Methods

### Mice

Six-to-eight-week-old female C57BL/6 mice were purchased from Charles River Laboratories France (Saint Germain Nuelles, France). *Rag2*^-/-^ (B6.129-Rag2tm1Fwa), Red5 mice (B6(C)-Il5tm1.1(icre)Lky/J)n, Rag2^-/-^γc^-/-^ (C;129S4-Rag2tm1.1Flv Il2rgtm1.1Flv/J) on a C57BL/6 J were bred in our animal facility. ROSA26-YFP mice (B6.129X1-*Gt(ROSA)26Sor^tm1(EYFP)Cos^*/J; 006148) were purchased from The Jackson Laboratory through Charles Rivers Laboratory France. All mice were maintained in specific-pathogen-free animal facility at IPBS and all experiments were conducted in strict accordance with French laws and regulations in compliance with the European Community council directive 68/609/EEC guidelines and its implementation in France under procedures approved by the French Ministry of Research and the FRBT (C2EA-01) animal care committee (APAFIS #1269, #3873, #10546, #16529 and #17384).

### *Mtb* culture, immunization & mouse infections

The laboratory strain of *Mtb*, H37Rv, was grown at 37°C in Middlebrook 7H9 medium (Difco) supplemented with 10% albumin-dextrose-catalase (ADC, Difco) and 0.05% Tyloxapol (Sigma), or on Middlebrook 7H11 agar medium (Difco) supplemented with 10% oleic acid-albumin-dextrose-catalase (OADC, Difco). Six- to eight-week-old mice were anesthetized with a cocktail of ketamine (60 mg/kg, Merial) and xylasine (10 mg/kg, Bayer) and infected intranasally (i.n.) with 1000 CFUs of mycobacteria in 25 μL of PBS containing 0.01% Tween 80. For immunization, C57BL/B6 mice were immunized i.n. with 5.10^5^ CFU of BCG (Danish), and were challenged 60 days post-vaccination with H37Rv as previously described. All experiments using *Mtb* were performed in appropriate biosafety level 3 (BSL3) laboratory and animal facility.

### *In vivo* treatments

2-DG (1g/kg, Sigma) was injected every other day starting from the day of infection and until completion of the experiment. For glucose supplementation, mice were treated with drinking water containing 30% (w/v) glucose (started 1 week before infection until sacrifice).

### Adoptive transfer experiments

For the adoptive transfer of total lung ILC2, *in vitro* cultured of ILC2 were harvested after 7 days of culture and 5×10^5^ to 2×10^6^ cells were transferred i.v. in mice anesthetized with isoflurane one day before *Mtb* infection in Rag2^-/-^γc^-/-^. For the adoptive transfer of the IL-18Rα^-^ or IL-18Rα^+^ ILC2 subsets, both subsets were FACS-sorted and cultured *in vitro* for 7 days in complete RPMI supplemented with 10% FBS. At the end of the culture, cells were harvested and 1.10^5^ cells were transferred i.v. in Rag2^-/-^γc^-/-^ mice anesthetized with isoflurane, one day before i.n. *Mtb* infection. For ILC1-like transfer, 1×10^4^ purified ILC1-like were directly transferred via intratracheal (i.t.) route in mice anesthetized with isoflurane one day before *Mtb* infection in Rag2^-/-^γc^-/-^.

### Lung harvest

Mice were sacrificed any cervical dislocation under isoflurane anesthesia and lungs were harvested aseptically, homogenized using a gentleMACS dissociator (C Tubes, Miltenyi) in HBSS (Difco), and incubated with DNAse I (0.1 mg/mL, Roche) and collagenase D (2 mg/mL, Roche) during 30 min at 37°C under 5% CO2. When indicated, mice received an i.v. injection of labeled anti-CD45 mAb (5μg) 5 minutes before sacrifice to discriminate between parenchymal and intravascular cells in subsequent flow cytometry analyses. Lung homogenates were filtered on 40 μm cell strainers and centrifuged at 329 × g during 5 min. Supernatants were conserved for cytokine content analysis. A part of the cellular pellet was conserved in TRIzol reagent for cellular RNA analysis. Bacterial loads (colony forming units) were determined by plating serial dilutions of the lung homogenates onto 7H10 solid medium (Difco) supplemented with 10% oleic acid-albumin-dextrose-catalase (OADC, Difco). The plates were incubated at 37°C for 3 weeks before bacterial CFUs scoring. In the remaining fraction, red blood cells were lysed in 150 mM NH4Cl, 10 mM KHCO_3_, 0.1 mM EDTA (pH 7.2) for immunological staining.

### *In situ* expansion of ILC

To expand ILC2, C57BL/6 or *Rag2*^-/-^ mice were treated intranasally (i.n.) with 100 ng of recombinant IL-33 (Biolegend) each day for 5 consecutive days. For the cytokine-based plasticity model, C57BL/6 or *Rag2*^-/-^ mice were treated i.n. with different combinations of cytokines specified in figures legends at day 1, 3, 5, 8 and sacrificed at day 9: 100 ng of IL-12 (R&D), IL-18 (R&D), IL-33 (Biolegend) or 20 μg of NMU (US Biological) per mouse and per instillation. For the Seahorse assays we elicited ILC2 with 0.5 mg IL-33, three doses i.p. over 10 days. Sorted ILC2 from lung were then cultured in presence of IL-7 and IL-2 (50ng/ml) for 7 days before addition of DMOG.

### Flow cytometry

To identify mouse ILCs, single-cell suspensions were stained with mAb for known lineages and with mAb discriminating ILC subsets. mAbs for known lineages included CD3 (17A2, Biolegend), CD4 (RM4-5, Biolegend), CD8a (53-6.7, Biolegend), TCRαβ (H57-597, Biolegend), TCRγδ, (GL3, Biolegend) CD11b (M1/70, Biolegend), CD11c (N418, Biolegend), F4/80 (BM8, Biolegend), Ly6G (1A8, Biolegend), TER119 (TER-119, Biolegend), FcεRIa (MAR-1, Biolegend), CD19 (1D3/CD19, Biolegend), B220 (RA3-6B2, Biolegend), and CD49b (DX5, Biolegend). mAbs discriminating ILC subsets included CD45.2 (104, BD), CD90.2 (30-H12, Biolegend), CD127 (A7R34, eBioscience), NK1.1 (PK136, BD Biosciences), IL-18Rα (P3TUNYA, eBioscience), ST2 (RMST2-2, eBioscience), CD226 (10E5, Biolegend), and CD49a (Ha31/8), NKp46 (29A1.4). mAbs for intracellular staining included GATA3 (L50-823, BD Biosciences), T-bet (4B10, eBiosciences), RORγt (Q31-378, BD Biosciences), TCF-1 (S33-966, BD), Arg1 (A1exF5, BD Biosciences), Ki-67 (SolA15, eBiosciences), Eomes (Dan11mag, eBiosciences, and HIF1α (D1S7W, Cell Signaling). After extracellular staining, cells were fixed and permeabilized (Foxp3 staining kit, eBiosciences) for intracellular staining. Samples from Biosafety Level 3 were inactivated for 2 hours at RT with 4% paraformaldehyde (ThermoFisher Scientific) after extracellular and intracellular staining.

Live/Dead fixable blue (eBiosciences) and mouse FcBlock (BD Biosciences) were used for all flow cytometry experiments. Cell staining was analyzed using LSR Fortessa flow cytometers (BD) and FlowJo software (v10). Cells were first gated in singlets (FSC-H vs. FSC-W and SSC-H vs. SSC-W) and live cells before further analyses.

### Intracellular cytokines staining

For intracellular cytokines staining of ILCs, single-cell suspensions from lung were incubated at 37°C with Brefeldin A in association or not with PMA (50 ng/ml, Sigma)/Ionomycine (500 ng/ml, Sigma) or 50 ng/ml of IL-12 and IL-18 for 4 hours before being surface stained, fixed and permeabilized (Foxp3 staining kit, eBiosciences). mAbs for cytokines staining included IFN-γ (XMG1.2, Biolegend), IL-17A (TC11-18H10, BD Biosciences), IL-5 (TRFK5, BD Biosciences), and IL-13 (eBio13A, eBiosciences) For HIF1α staining of ILCs, single-cell suspensions from lung were incubated at 37°C with DMOG (500μM) for 3 hours before being surface stained, fixed and permeabilized (Foxp3 staining kit, eBiosciences). In order to block glycolysis during *ex vivo* stimulation, cells were incubated in the presence of 10mM 2-DG (Sigma). *Mtb was* inactivated by incubation in PFA 4% for 2 hours at room temperature.

### ILC enrichment and cell-sorting

Lung ILCs were enriched from lung single-cell suspensions by using the EasySep^TM^ Mouse ILC2 Enrichment Kit (StemCell). After enrichment, cells were stained with lineage mAb (CD3, CD4, CD8α, TCRαβ, TCRγδ, CD19, B220, CD11b, CD11c, F4/80, TER119, FcεRIa, CD49b, Ly6G) and ILC markers (CD90.2, CD45.2, NK1.1, ST2, IL-18Rα, CD49a). ILC2 were purified as Lin^-^CD45.2^+^CD90.2^+^NK1.1^-^ST2^+^IL-18Rα^+/-^ ILC1-like were purified as Lin^-^CD45.2^+^CD90.2^+^NK1.1^-^ST2^-^CD49a^+^IL-18Rα^+^. Cells were sorted using a FACSAria Fusion cytometer (BD, France).

### *In vitro* culture of ILC2

Cell sorted ILC2 were incubated in 6-well plates at a density of 300,000 cells per ml for 4 days with IL-2 (25 ng/ml, R&D) and IL-7 (25 ng/ml, R&D) in RPMI (Difco) supplemented with 10 % FBS. After 4 days of culture, ILC2 were harvested for adoptive transfer or incubated with DMOG (Sigma). For DMOG experiment, half of the medium was removed and replaced with fresh medium containing IL-2 (25 ng/ml) and IL-7 (25 ng/ml) with or without DMOG (500 μM) for 3 more days. Cell sorted ILC2 were incubated in 6-well plates at a density of 300,000 cells per ml for 4 days with IL-2 (25 ng/ml, R&D) and IL-7 (25 ng/ml, R&D) in RPMI (Difco) supplemented with 10 % FBS. After 4 days of culture, ILC2 were harvested for adoptive transfer or incubated with DMOG (Sigma). For DMOG experiment, half of the medium was removed and replaced with fresh medium containing IL-2 (25 ng/ml) and IL-7 (25 ng/ml) with or without DMOG (500 μg/ml) for 3 more days. For the Seahorse assays, sorted ILC2 from lung were cultured in presence of IL-7 and IL-2 (50ng/ml) for 7 days before addition of DMOG.

### SCENITH assay

SCENITH experiments were performed as previously described (Argüello et al., 2020) using the SCENITH kit containing all reagents and anti-puromycin antibodies (requested from www.scenith.com/try-it). Briefly, lung cell suspensions were stimulated for 15 minutes at 37°C in the presence of the indicated inhibitors of various metabolic pathways then incubated for 30 minutes with puromycin at 37°C. At the end of the incubation, puromycin was stained with fluorescent anti-puromycin antibodies (Clone R4743L-E8) by flow cytometry and the impact of the various metabolic inhibitors was quantitated as described (Argüello et al., 2020).

### Seahorse experiments

1.5 to 2×10^5^ FACS sorted lung ILC2s per well were rested in a 96-well plate in Glutamax RPMI (supplemented with 10% fetal bovine serum, non-essential amino acids, 1 mM sodium pyruvate, 85 μM 2-mercapto-ethanol and 100 U/ml penicillin-streptomycin) containing 25 ng/ml IL-7. After 24h cells were split and rested in fresh IL-7 containing media for another 3 days. Subsequently, cells were cultured in fresh medium containing 25 ng/ml IL-7 and 20 ng/ml of IL-2 in the presence or absence of 0.5 mM DMOG for a further 72 hours. To prepare for extracellular flux analysis cells were then washed thoroughly in XF medium (modified DMEM) and adhered to the Seahorse plate using 22.4 μg/ml Cell-Tak (Corning).

For glycolytic stress test, cells were plated at a density of 2×10^5^ cells/well in XF medium supplemented with 2 mM glutamine. Cells were incubated for 30-60 min at 37°C and ECAR was measured under basal conditions, and in response to 10 mM glucose, 2 μM oligomycin and 50 mM 2-DG. For the mitochondrial stress test, cells were plated at a density of 1.5×10^5^ cells/well in XF medium supplemented with 2 mM glutamine, 1 mM sodium pyruvate and 25 mM glucose. Cells were incubated for 30-60 min at 37°C. OCAR was measured under basal conditions, after injection of 2 μM oligomycin, 1.5 μM FCCP and 100 nM rotenone + 1 μM antimycin A. Extracellular flux assays were done using a 96-well extracellular flux analyzer XFe-96 (Seahorse Bioscience).

Normalization by protein was used to correct for potential differences in seeding densities across wells. Protein measurement was performed using the Pierce BCA protein assay according to the manufacturer instructions.

### Quantitative RT-PCR analysis of transcripts

RNA from lungs homogenates was extracted using TRIzol reagent (Ambion) and RNeasy spin columns according to manufacturer’s instructions (RNeasy kit, Qiagen). RNA was reverse transcribed into cDNA using M-MLV Reverse transcriptase (Invitrogen). RT-qPCR was performed using gene-targeted primers (Supplementary Table 1) as described above. Values were normalized using the housekeeping beta-actin gene (*Actb*) and expressed as a fold change. RNA from ILC2 culture were extracted using RLT (Qiagen) and RNA were reverse transcribed as previously described (Troegeler et al., 2017).

### Statistical analyses

Statistical analyses were performed using GraphPad Prism 9 software. Agostino and Pearson normality tests were performed to determine whether data followed a normal distribution. Unpaired *t*-test (for normal data) or Mann-Whitney (for non-normal data) were performed when two samples were compared; ANOVA (for normal data) or Kruskal-Wallis (for non-normal data) tests were performed when more than two samples were compared. For all analyses, * indicates P < 0.05, ** indicates P < 0.01, *** indicates P < 0.001, and **** indicates P < 0.0001.

## Acknowledgements

We thank Yasmine Belkaid (NIH/NIAID, Bethesda) for critical review of the manuscript. We acknowledge Emmanuelle Näser (Genotoul TRI-IPBS platform, Toulouse) for flow cytometry and cell-sorting, Flavie Moreau, Céline Berrone and Aline Tridon (Genotoul Anexplo-IPBS platform, Toulouse), and Sylvie Appolinaire and Celine Berraud (CREFRE US006, Toulouse) for mouse care and maintenance in conventional and BSL3 facilities. We thank Richard Locksley (USCF, San Francisco) for the kind gift of Red5 mice. We thank Geanncarlo Lugo-Villarino, Sophie Laffont (CPTP, Toulouse) and Andrea Pichler (CRCT, Toulouse) for helpful discussions. This work was supported by Centre de la Recherche Scientifique (CNRS), the University Toulouse III-Paul Sabatier, the French Ministry for Higher Education, Research and Innovation (fellowship to D.C.), the Fondation pour la Recherche Médicale (grant DEQ20160334902 to O.N.), the Bettencourt Schueller Foundation (grants Coup d’Élan pour la recherche française and Explore-TB to O.N.), MSDAVENIR (grant Fight-TB to O.N.), the Agence Nationale de la Recherche (grants ANR-18-CE15-0004-01 to D.H. and ANR-11-EQUIPEX-0003 to O.N.), and the European Commission (grant TBVAC2020 n°643381 to O.N.). M.R.H. is supported by a Royal Society and Wellcome Trust Sir Henry Dale fellowship (105644/Z/14/Z), a Lister Institute of Preventative Medicine Prize and a BBSRC Project grant (BB/T014482/1). The funders had no role in study design, data collection, and analysis, decision to publish, or preparation of the manuscript.

## Author contributions

D. C. and D.H. conceived and designed the study with input from O.N.; D.C., A.C., M.Z.K., E.B., T.S. and F.L. performed the experiments; E.L., J-P.G., and R.J.A contributed critical reagents and methods, D.C., G.E., J-C.G., R.J.A., M.R.H. and D.H. analyzed and interpreted the data; J-C.G., G.E., M.R.H., and Y.P., also provide important discussion for the project and critical feedback on the manuscript; D.C., O.N. and D.H. wrote the manuscript. All coauthors read, reviewed and approved the manuscript.

## Competing interests

The authors declare no competing interests

**Supplementary Figure 1.**
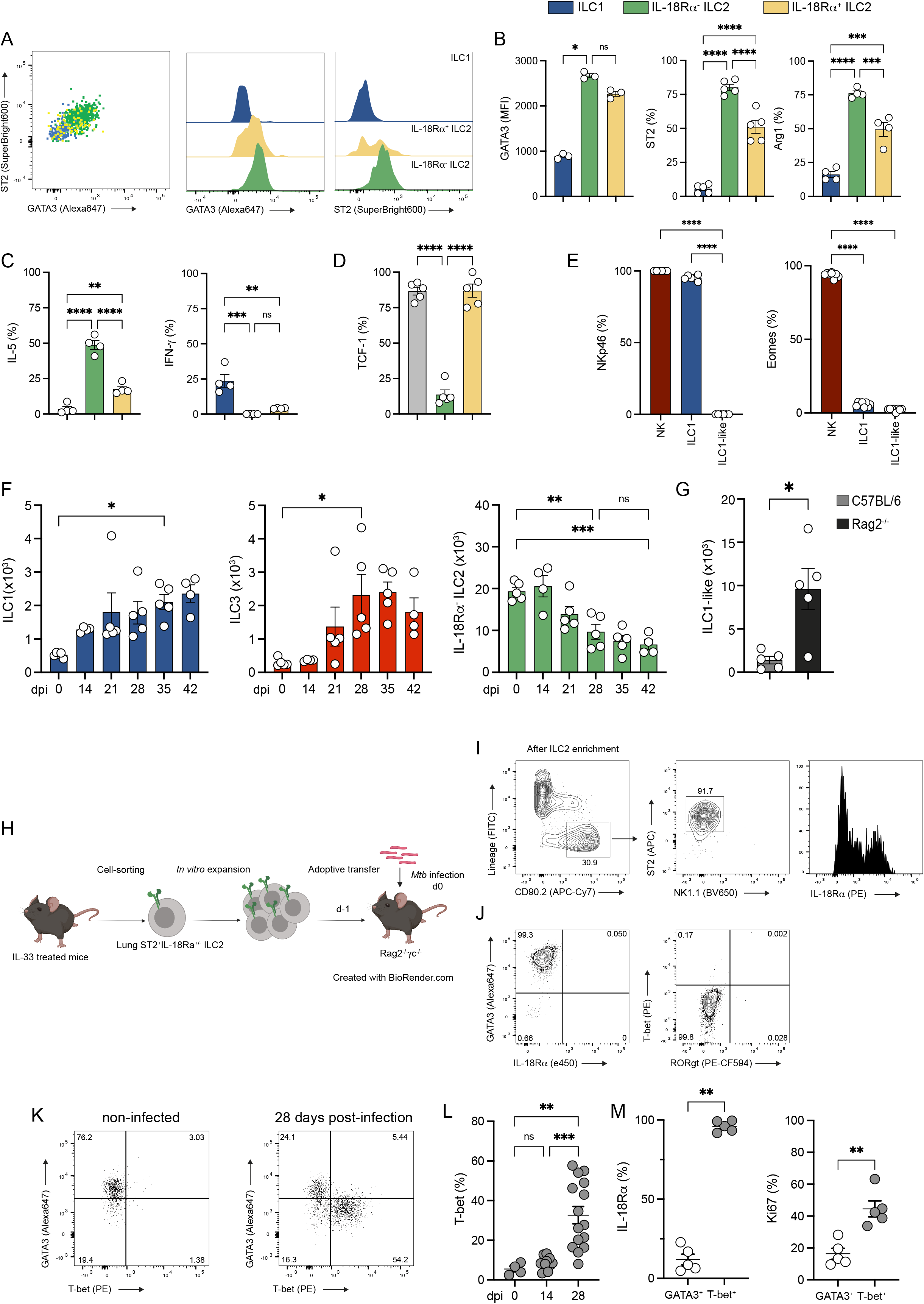
Dynamics of ILCs during *Mtb* infection in the mouse model. **(A)** Representative expression of GATA3 and ST2 on the indicated ILC subsets. **(B)** Expression of GATA3 (MFI), ST2 (%) and Arg1 (%) in ILC1 (blue), IL-18Rα^-^ ILC2 (green) and IL-18Rα^+^ ILC2 (yellow) at steady-state in the lung of C57BL/6 mice. **(C)** Expression of IL-5 (%) and IFN-γ (%) in the indicated ILC subsets after *ex vivo* stimulation with PMA/ionomycin in presence of Brefeldin A for 4h at steady-state in the lung of C57BL/6 mice **(D)** Percentages of TCF-1^+^ cells in ILC2P from bone marrow (grey) compared to IL-18Rα^-^ ILC2 (green) and IL-18Rα^+^ ILC2 (yellow) from the lungs of C57BL/6 mice at steady-state. (**E)** Percentages of NKp46^+^ (left) and Eomes^+^ (right) cells in ILC1-like cells compared to in NK cells and ILC1 at day 28 post- infection. **(F)** Absolute numbers of ILC1 (dark blue), ILC3 (red) and IL-18Rα^-^ ILC2 at the indicated days after *Mtb* infection. Prior to sacrifice, mice were injected with fluorescent anti-CD45.2 to distinguish vascular and parenchymal cells. ILC1, ILC3, IL- 18Rα^-^ ILC2 and IL-18Rα^+^ ILC2 have been gated on lung-resident cells. **(G)** Absolute number of ILC1-like cells at day 28 post-infection in C57BL/5 (grey) *vs.* Rag2^-/-^ (black) mice. **(H)** Schematic representation of the *in vivo* expansion of ILC2 in C57BL/6 or *Rag2*^-/-^ mice treated with IL-33, cell-sorting, *in vitro* culture and adoptive transfer of ILC2 in *Rag2*^-/-^*Il2rg*^-/-^ one day before infection with *Mtb*. **(I)** Gating strategy to purify ILC2 based on the expression of ST2 (left two graphs) and purity of ILC2 after cell-sorting (right). **(J)** Phenotype of ILC2 after *in vitro* culture and before adoptive transfer. **(K)** A representative dot-plot of GATA3 and T-bet expression in Lin^-^CD45.2^+^CD90.2^+^ cells isolated from *Rag2*^-/-^*Il2rg*^-/-^ mice adoptively transferred with purified ILC2 then left uninfected (right) or infected with *Mtb* (left). **(L)** Percentages of T-bet expressing ILC at different days post-infection. **(M)** Expression of IL-18Rα (%) and Ki67 (%) in transferred GATA3^+^ (white dots) *vs.* T-bet^+^ (grey dots) ILC at day 28 post-infection in *Rag2*^-/-^*Il2rg*^-/-^ mice. Each symbol represents an individual mouse. Statistical analysis was performed using one-way ANOVA test **(B-F, L)** or Mann-Whitney test **(G, M)** (*, p<0.05; **, P<0.01; ***, p<0.001; ****, p<0.0001). Graphs depict data as mean (± s.e.m). Data are representative of five **(F)**, three **(B-C, K-M)** and two **(D, E, G)** independent experiments.

**Supplementary Figure 2.**
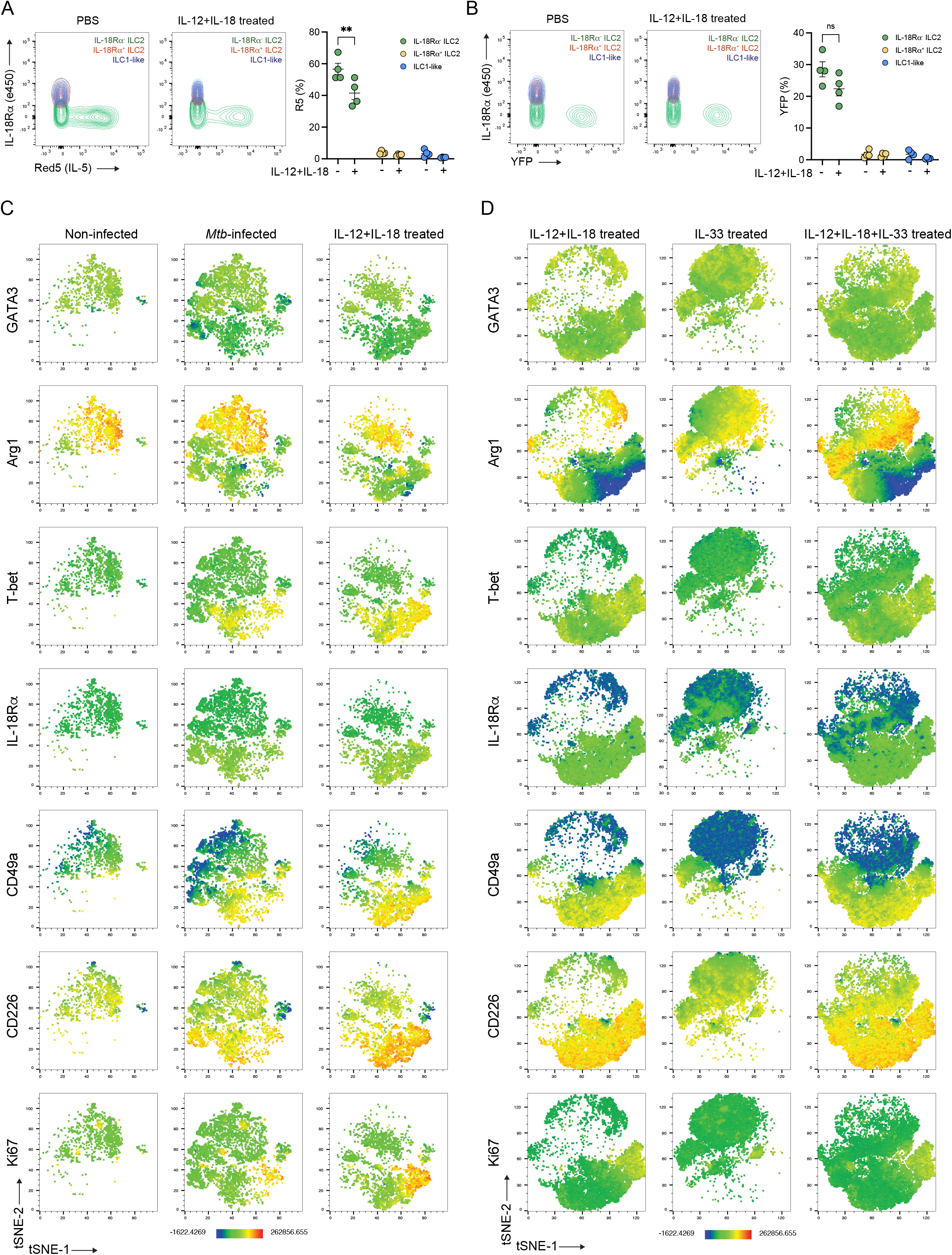
Combination of IL-12, IL-18 and IL-33 induces a mixed ILC1 and ILC2 phenotype on IL-18Rα^+^ ILC2. **(A)** Representative dot-plot showing IL- 5 and IL-18Rα expression among IL-18Rα^-^ ILC2 (green), IL-18Rα^+^ ILC2 (red), and ILC1-like (blue) in PBS *vs*. IL-12+IL-18 treated IL-5^Cre^ROSA26^YFP^ mic (left) and quantification (right). **(B)** Representative dot-plot showing YFP and IL-18Rα expression among IL-18Rα^-^ ILC2 (green), IL-18Rα^+^ ILC2 (red), and ILC1-like (blue) in PBS *vs*. IL-12+IL-18 treated IL-5^Cre^ROSA26^YFP^ mic (left) and quantification (right). **(C)** Unsupervised t-SNE representation of the expression of different markers (GATA3, Arg1, T-bet, IL-18Rα, CD49a, CD226 and Ki67) on Lin^-^CD45.2^+^CD90.2^+^NK1.1^-^RORγt^-^ in non-infected *vs*. *Mtb*-infected (28 dpi) *vs.* IL-12+IL-18 treated Rag2^-/-^ mice. **(D)** As in A) except that the analysis was performed on Lin^-^CD45.2^+^CD90.2^+^NK1.1^-^RORγt^-^ in IL-12+IL-18 *vs*. IL-33 *vs.* IL-12+IL-18 treated Rag2^-/-^ mice. Statistical analysis was performed two-way ANOVA test **(A-B)** (*, p<0.05; **, P<0.01; ***, p<0.001; ****, p<0.0001). Graphs depict data as mean (± s.e.m). Data are representative of two **(A-B)** independent experiments.

**Supplementary Figure 3.**
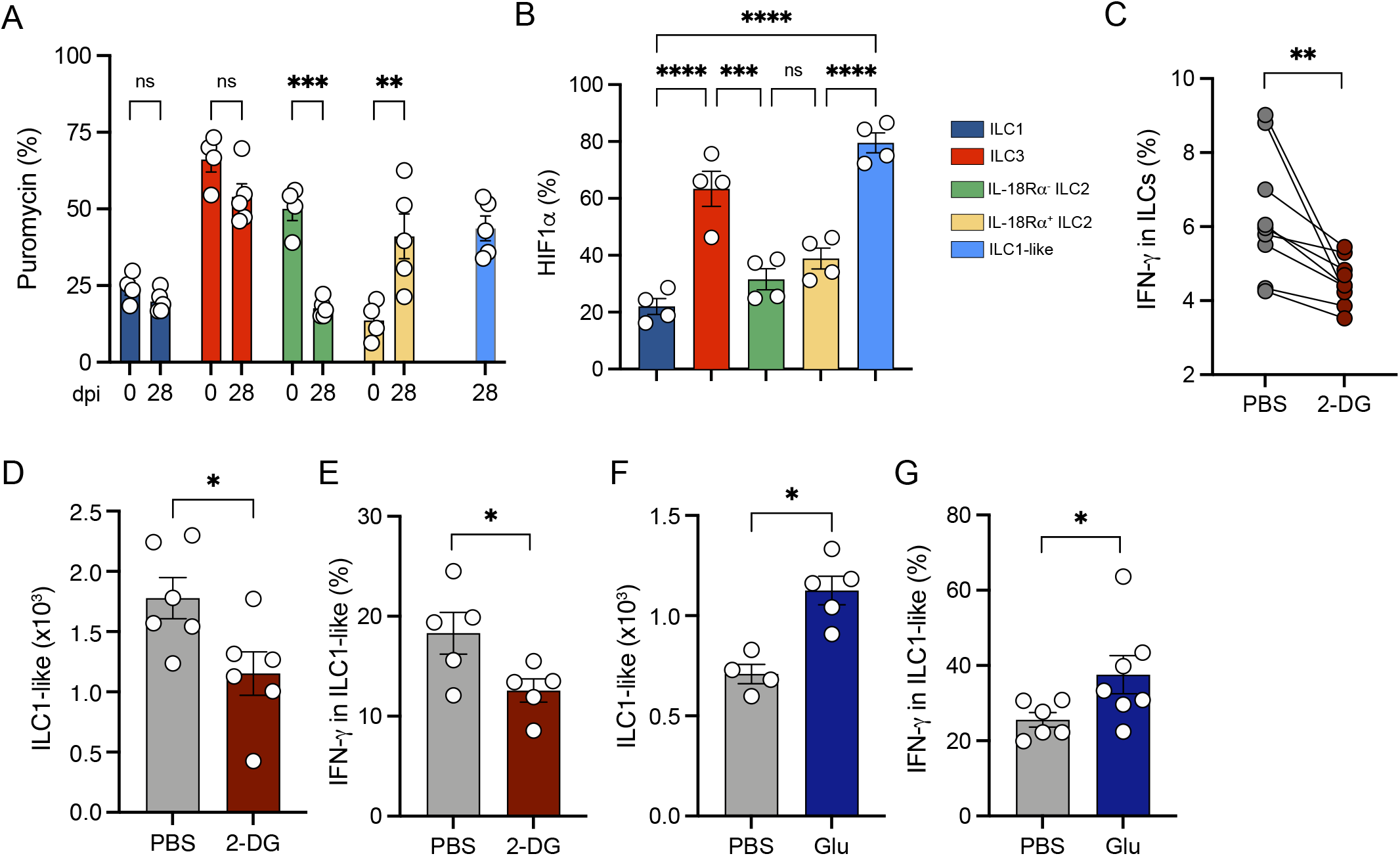
Glycolysis regulates ILC1-like cell differentiation during *Mtb* infection. **(A)** Percentage of puromycin positive cells in ILC1 (dark blue), ILC3 (red), IL-18Rα^-^ ILC2 (green), IL-18Rα^+^ ILC2 (yellow) and ILC1-like cells (blue) in non-infected *vs. Mtb*-infected Rag2^-/-^ mice. **(B)** HIF1α expression in the indicated ILC subsets at day 28 post-infection **(C)** Expression of IFN-γ in total ILCs after *ex vivo* stimulation with IL-12+IL-18 in the presence or absence of 2-DG**. (D)** Absolute numbers of ILC1-like cells in *Rag2*^-/-^ mice treated or not with 2-DG during *Mtb* infection at day 28 post-infection. **(E)** Percentages of IFNγ-producing cells among ILC1-like cells after *ex vivo* stimulation with PMA/ionomycin in the presence of brefeldin A for 4h from PBS *vs.* 2-DG treated mice. **(F)** As in **(D)** except that mice treated with 30% glucose in their drinking water. **(G)** As in **(E)** except that mice were treated or not with 30% glucose in their drinking water. Each symbol represents an individual mouse and statistical analysis was performed using two-way ANOVA **(A)**, one-way ANOVA **(B)**, Wilcoxon **(C)** and Mann-Whitney **(D-G)** tests (*, p<0.05; **, P<0.01; ***, p<0.001; ****, p<0.0001). Graphs depict data as mean (± s.e.m) from three **(D-E)** or two **(A-C, F-G)** independent experiments.

**Supplementary Figure 4.**
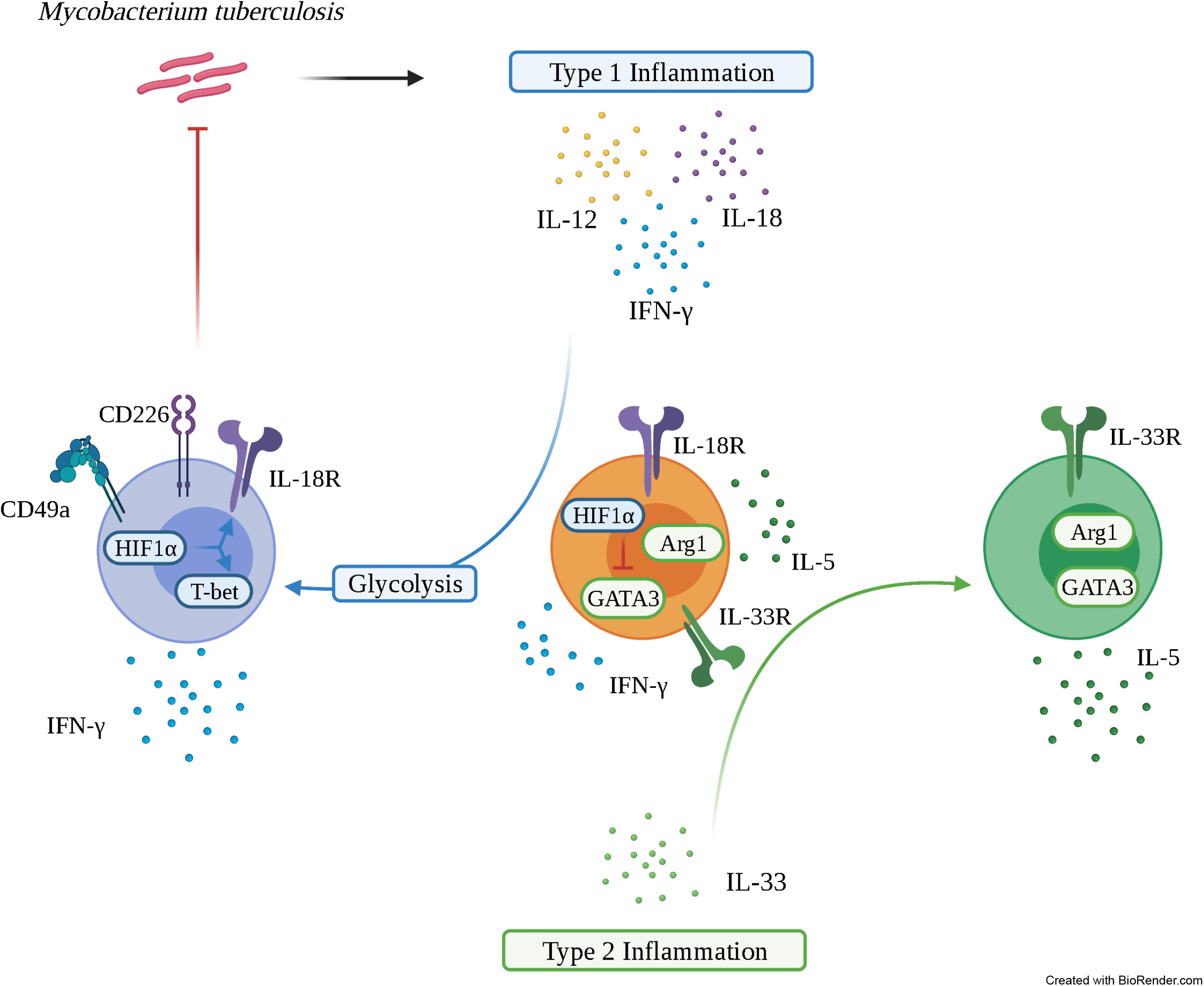
Metabolic regulation of IL-18Rα^+^ ILC2 differentiation into ILC1-like cells during *Mycobacterium tuberculosis* infection. *Mtb* infection results in the establishment of a type 1 inflammation. Type 1 cytokines, IL-12, IL-18 and IFN-γ (upper part) act on a rare, lung immature IL-18Rα-expressing ILC2 subset and triggers a glycolysis-involving metabolic reprogramming leading to its differentiation into ILC1-like cells IL18Rα^+^, CD49a^+^CD226^+^HIF-1α^+^, T-bet^+^, IFN-γ-producing) endowed with a protective potential against *Mtb*. In contrast, type 2 cytokines such as IL-33 (lower part) acts on this immature IL-18Rα-expressing ILC2 subset to drive its maturation toward mature IL-5 producing ILC2.

## Notes

### Competing Interest Statement

The authors have declared no competing interest.

## References

1. Ardain, A., Domingo-Gonzalez, R., Das, S., Kazer, S.W., Howard, N.C., Singh, A., Ahmed, M., Nhamoyebonde, S., Rangel-Moreno, J., Ogongo, P., Lu, L., Ramsuran, D., de la Luz Garcia-Hernandez, M., K. Ulland, T., Darby, M., Park, E., Karim, F., Melocchi, L., Madansein, R., Dullabh, K.J., Dunlap, M., Marin-Agudelo, N., Ebihara, T., Ndung’u, T., Kaushal, D., Pym, A.S., Kolls, J.K., Steyn, A., Zúñiga, J., Horsnell, W., Yokoyama, W.M., Shalek, A.K., Kløverpris, H.N., Colonna, M., Leslie, A., Khader, S.A., 2019. Group 3 innate lymphoid cells mediate early protective immunity against tuberculosis. Nature 570, 528–532. https://doi.org/10.1038/s41586-019-1276-2

2. Argüello, R.J., Combes, A.J., Char, R., Gigan, J.-P., Baaziz, A.I., Bousiquot, E., Camosseto, V., Samad, B., Tsui, J., Yan, P., Boissonneau, S., Figarella-Branger, D., Gatti, E., Tabouret, E., Krummel, M.F., Pierre, P., 2020. SCENITH: A Flow Cytometry-Based Method to Functionally Profile Energy Metabolism with Single-Cell Resolution. Cell Metab. 32, 1063–1075.e7. https://doi.org/10.1016/j.cmet.2020.11.007

3. Bal, S.M., Golebski, K., Spits, H., 2020. Plasticity of innate lymphoid cell subsets. Nat. Rev. Immunol. 1–14. https://doi.org/10.1038/s41577-020-0282-9

4. Bando, J.K., Nussbaum, J.C., Liang, H.-E., Locksley, R.M., 2013. Type 2 innate lymphoid cells constitutively express arginase-I in the naïve and inflamed lung. J. Leukoc. Biol. 94, 877–884. https://doi.org/10.1189/jlb.0213084

5. Cardoso, V., Chesné, J., Ribeiro, H., García-Cassani, B., Carvalho, T., Bouchery, T., Shah, K., Barbosa-Morais, N.L., Harris, N., Veiga-Fernandes, H., 2017. Neuronal regulation of type 2 innate lymphoid cells via neuromedin U. Nature 549, 277–281. https://doi.org/10.1038/nature23469

6. Chiossone, L., Dumas, P.-Y., Vienne, M., Vivier, E., 2018. Natural killer cells and other innate lymphoid cells in cancer. Nat. Rev. Immunol. 18, 671–688. https://doi.org/10.1038/s41577-018-0061-z

7. Di Luccia, B., Gilfillan, S., Cella, M., Colonna, M., Huang, S.C.-C., 2019. ILC3s integrate glycolysis and mitochondrial production of reactive oxygen species to fulfill activation demands. J. Exp. Med. 216, 2231–2241. https://doi.org/10.1084/jem.20180549

8. Fernández-García, M., Rey-Stolle, F., Boccard, J., Reddy, V.P., García, A., Cumming, B.M., Steyn, A.J.C., Rudaz, S., Barbas, C., 2020. Comprehensive Examination of the Mouse Lung Metabolome Following Mycobacterium tuberculosis Infection Using a Multiplatform Mass Spectrometry Approach. J. Proteome Res. 19, 2053–2070. https://doi.org/10.1021/acs.jproteome.9b00868

9. Ghaedi, M., Shen, Z.Y., Orangi, M., Martinez-Gonzalez, I., Wei, L., Lu, X., Das, A., Heravi-Moussavi, A., Marra, M.A., Bhandoola, A., Takei, F., 2020. Single-cell analysis of RORα tracer mouse lung reveals ILC progenitors and effector ILC2 subsets. J. Exp. Med. 217. https://doi.org/10.1084/jem.20182293

10. Gopal, R., Monin, L., Slight, S., Uche, U., Blanchard, E., Fallert Junecko, B.A., Ramos-Payan, R., Stallings, C.L., Reinhart, T.A., Kolls, J.K., Kaushal, D., Nagarajan, U., Rangel-Moreno, J., Khader, S.A., 2014. Unexpected role for IL-17 in protective immunity against hypervirulent Mycobacterium tuberculosis HN878 infection. PLoS Pathog. 10, e1004099. https://doi.org/10.1371/journal.ppat.1004099

11. Gury-BenAri, M., Thaiss, C.A., Serafini, N., Winter, D.R., Giladi, A., Lara-Astiaso, D., Levy, M., Salame, T.M., Weiner, A., David, E., Shapiro, H., Dori-Bachash, M., Pevsner-Fischer, M., Lorenzo-Vivas, E., Keren-Shaul, H., Paul, F., Harmelin, A., Eberl, G., Itzkovitz, S., Tanay, A., Di Santo, J.P., Elinav, E., Amit, I., 2016. The Spectrum and Regulatory Landscape of Intestinal Innate Lymphoid Cells Are Shaped by the Microbiome. Cell 166, 1231–1246.e13. https://doi.org/10.1016/j.cell.2016.07.043

12. Heiden, M.G.V., Cantley, L.C., Thompson, C.B., 2009. Understanding the Warburg Effect: The Metabolic Requirements of Cell Proliferation. Science 324, 1029–1033. https://doi.org/10.1126/science.1160809

13. Joseph, A.M., Monticelli, L.A., Sonnenberg, G.F., 2018. Metabolic regulation of innate and adaptive lymphocyte effector responses. Immunol. Rev. 286, 137–147. https://doi.org/10.1111/imr.12703

14. Kinjo, Y., Kawakami, K., Uezu, K., Yara, S., Miyagi, K., Koguchi, Y., Hoshino, T., Okamoto, M., Kawase, Y., Yokota, K., Yoshino, K., Takeda, K., Akira, S., Saito, A., 2002. Contribution of IL-18 to Th1 response and host defense against infection by Mycobacterium tuberculosis: a comparative study with IL-12p40. J. Immunol. Baltim. Md 1950 169, 323–329. https://doi.org/10.4049/jimmunol.169.1.323

15. Klose, C.S.N., Flach, M., Möhle, L., Rogell, L., Hoyler, T., Ebert, K., Fabiunke, C., Pfeifer, D., Sexl, V., Fonseca-Pereira, D., Domingues, R.G., Veiga-Fernandes, H., Arnold, S.J., Busslinger, M., Dunay, I.R., Tanriver, Y., Diefenbach, A., 2014. Differentiation of Type 1 ILCs from a Common Progenitor to All Helper-like Innate Lymphoid Cell Lineages. Cell 157, 340–356. https://doi.org/10.1016/j.cell.2014.03.030

16. Li, Q., Li, D., Zhang, X., Wan, Q., Zhang, W., Zheng, M., Zou, L., Elly, C., Lee, J.H., Liu, Y.-C., 2018. E3 Ligase VHL Promotes Group 2 Innate Lymphoid Cell Maturation and Function via Glycolysis Inhibition and Induction of Interleukin-33 Receptor. Immunity 48, 258–270.e5. https://doi.org/10.1016/j.immuni.2017.12.013

17. Lim, A.I., Di Santo, J.P., 2019. ILC-poiesis: Ensuring tissue ILC differentiation at the right place and time. Eur. J. Immunol. 49, 11–18. https://doi.org/10.1002/eji.201747294

18. Lim, A.I., Li, Y., Lopez-Lastra, S., Stadhouders, R., Paul, F., Casrouge, A., Serafini, N., Puel, A., Bustamante, J., Surace, L., Masse-Ranson, G., David, E., Strick-Marchand, H., Bourhis, L.L., Cocchi, R., Topazio, D., Graziano, P., Muscarella, L.A., Rogge, L., Norel, X., Sallenave, J.-M., Allez, M., Graf, T., Hendriks, R.W., Casanova, J.-L., Amit, I., Yssel, H., Santo, J.P.D., 2017. Systemic Human ILC Precursors Provide a Substrate for Tissue ILC Differentiation. Cell 168, 1086–1100.e10. https://doi.org/10.1016/j.cell.2017.02.021

19. Manca, C., Tsenova, L., Bergtold, A., Freeman, S., Tovey, M., Musser, J.M., Barry, C.E., Freedman, V.H., Kaplan, G., 2001. Virulence of a Mycobacterium tuberculosis clinical isolate in mice is determined by failure to induce Th1 type immunity and is associated with induction of IFN-α/β. Proc. Natl. Acad. Sci. 98, 5752–5757. https://doi.org/10.1073/pnas.091096998

20. Meininger, I., Carrasco, A., Rao, A., Soini, T., Kokkinou, E., Mjösberg, J., 2020. Tissue-Specific Features of Innate Lymphoid Cells. Trends Immunol. 41, 902–917. https://doi.org/10.1016/j.it.2020.08.009

21. Monticelli, L.A., Buck, M.D., Flamar, A.-L., Saenz, S.A., Tait Wojno, E.D., Yudanin, N.A., Osborne, L.C., Hepworth, M.R., Tran, S.V., Rodewald, H.-R., Shah, H., Cross, J.R., Diamond, J.M., Cantu, E., Christie, J.D., Pearce, E.L., Artis, D., 2016. Arginase 1 is an innate lymphoid-cell-intrinsic metabolic checkpoint controlling type 2 inflammation. Nat. Immunol. 17, 656–665. https://doi.org/10.1038/ni.3421

22. Moro, K., Yamada, T., Tanabe, M., Takeuchi, T., Ikawa, T., Kawamoto, H., Furusawa, J., Ohtani, M., Fujii, H., Koyasu, S., 2010. Innate production of T H 2 cytokines by adipose tissue-associated c-Kit + Sca-1 + lymphoid cells. Nature 463, 540–544. https://doi.org/10.1038/nature08636

23. Neill, D.R., Wong, S.H., Bellosi, A., Flynn, R.J., Daly, M., Langford, T.K.A., Bucks, C., Kane, C.M., Fallon, P.G., Pannell, R., Jolin, H.E., McKenzie, A.N.J., 2010. Nuocytes represent a new innate effector leukocyte that mediates type-2 immunity. Nature 464, 1367–1370. https://doi.org/10.1038/nature08900

24. Nussbaum, J.C., Van Dyken, S.J., von Moltke, J., Cheng, L.E., Mohapatra, A., Molofsky, A.B., Thornton, E.E., Krummel, M.F., Chawla, A., Liang, H.-E., Locksley, R.M., 2013. Type 2 innate lymphoid cells control eosinophil homeostasis. Nature 502, 245–248. https://doi.org/10.1038/nature12526

25. O’Garra, A., Redford, P.S., McNab, F.W., Bloom, C.I., Wilkinson, R.J., Berry, M.P.R., 2013. The Immune Response in Tuberculosis. Annu. Rev. Immunol. 31, 475–527. https://doi.org/10.1146/annurev-immunol-032712-095939

26. Palazon, A., Goldrath, A.W., Nizet, V., Johnson, R.S., 2014. HIF Transcription Factors, Inflammation, and Immunity. Immunity 41, 518–528. https://doi.org/10.1016/j.immuni.2014.09.008

27. Perdomo, C., Zedler, U., Kühl, A.A., Lozza, L., Saikali, P., Sander, L.E., Vogelzang, A., Kaufmann, S.H.E., Kupz, A., 2016. Mucosal BCG Vaccination Induces Protective Lung-Resident Memory T Cell Populations against Tuberculosis. mBio 7. https://doi.org/10.1128/mBio.01686-16

28. Price, A.E., Liang, H.-E., Sullivan, B.M., Reinhardt, R.L., Eisley, C.J., Erle, D.J., Locksley, R.M., 2010. Systemically dispersed innate IL-13–expressing cells in type 2 immunity. Proc. Natl. Acad. Sci. 107, 11489–11494. https://doi.org/10.1073/pnas.1003988107

29. Ricardo-Gonzalez, R.R., Van Dyken, S.J., Schneider, C., Lee, J., Nussbaum, J.C., Liang, H.-E., Vaka, D., Eckalbar, W.L., Molofsky, A.B., Erle, D.J., Locksley, R.M., 2018. Tissue signals imprint ILC2 identity with anticipatory function. Nat. Immunol. 19, 1093–1099. https://doi.org/10.1038/s41590-018-0201-4

30. Saluzzo, S., Gorki, A.-D., Rana, B.M.J., Martins, R., Scanlon, S., Starkl, P., Lakovits, K., Hladik, A., Korosec, A., Sharif, O., Warszawska, J.M., Jolin, H., Mesteri, I., McKenzie, A.N.J., Knapp, S., 2017. First-Breath-Induced Type 2 Pathways Shape the Lung Immune Environment. Cell Rep. 18, 1893–1905. https://doi.org/10.1016/j.celrep.2017.01.071

31. Schneider, C., Lee, J., Koga, S., Ricardo-Gonzalez, R.R., Nussbaum, J.C., Smith, L.K., Villeda, S.A., Liang, H.-E., Locksley, R.M., 2019. Tissue-Resident Group 2 Innate Lymphoid Cells Differentiate by Layered Ontogeny and In Situ Perinatal Priming. Immunity 50, 1425–1438.e5. https://doi.org/10.1016/j.immuni.2019.04.019

32. Shi, L., Salamon, H., Eugenin, E.A., Pine, R., Cooper, A., Gennaro, M.L., 2015. Infection with Mycobacterium tuberculosis induces the Warburg effect in mouse lungs. Sci. Rep. 5, 1–13. https://doi.org/10.1038/srep18176

33. Shih, H.-Y., Sciumè, G., Mikami, Y., Guo, L., Sun, H.-W., Brooks, S.R., Urban, J.F., Davis, F.P., Kanno, Y., O’Shea, J.J., 2016. Developmental Acquisition of Regulomes Underlies Innate Lymphoid Cell Functionality. Cell 165, 1120–1133. https://doi.org/10.1016/j.cell.2016.04.029

34. Silver, J.S., Kearley, J., Copenhaver, A.M., Sanden, C., Mori, M., Yu, L., Pritchard, G.H., Berlin, A.A., Hunter, C.A., Bowler, R., Erjefalt, J.S., Kolbeck, R., Humbles, A.A., 2016. Inflammatory triggers associated with exacerbations of COPD orchestrate plasticity of group 2 innate lymphoid cells in the lungs. Nat. Immunol. 17, 626–635. https://doi.org/10.1038/ni.3443

35. Svedberg, F.R., Brown, S.L., Krauss, M.Z., Campbell, L., Sharpe, C., Clausen, M., Howell, G.J., Clark, H., Madsen, J., Evans, C.M., Sutherland, T.E., Ivens, A.C., Thornton, D.J., Grencis, R.K., Hussell, T., Cunoosamy, D.M., Cook, P.C., MacDonald, A.S., 2019. The lung environment controls alveolar macrophage metabolism and responsiveness in type 2 inflammation. Nat. Immunol. 20, 571–580. https://doi.org/10.1038/s41590-019-0352-y

36. Troegeler, A., Mercier, I., Cougoule, C., Pietretti, D., Colom, A., Duval, C., Vu Manh, T.-P., Capilla, F., Poincloux, R., Pingris, K., Nigou, J., Rademann, J., Dalod, M., Verreck, F.A.W., Al Saati, T., Lugo-Villarino, G., Lepenies, B., Hudrisier, D., Neyrolles, O., 2017. C-type lectin receptor DCIR modulates immunity to tuberculosis by sustaining type I interferon signaling in dendritic cells. Proc. Natl. Acad. Sci. U. S. A. 114, E540–E549. https://doi.org/10.1073/pnas.1613254114

37. Urdahl, K., Shafiani, S., Ernst, J., 2011. Initiation and regulation of T-cell responses in tuberculosis. Mucosal Immunol. 4, 288–293. https://doi.org/10.1038/mi.2011.10

38. Vivier, E., Artis, D., Colonna, M., Diefenbach, A., Santo, J.P.D., Eberl, G., Koyasu, S., Locksley, R.M., McKenzie, A.N.J., Mebius, R.E., Powrie, F., Spits, H., 2018. Innate Lymphoid Cells: 10 Years On. Cell 174, 1054–1066. https://doi.org/10.1016/j.cell.2018.07.017

39. Vivier, E., van de Pavert, S.A., Cooper, M.D., Belz, G.T., 2016. The evolution of innate lymphoid cells. Nat. Immunol. 17, 790–794. https://doi.org/10.1038/ni.3459

40. Wallrapp, A., Riesenfeld, S.J., Burkett, P.R., Abdulnour, R.-E.E., Nyman, J., Dionne, D., Hofree, M., Cuoco, M.S., Rodman, C., Farouq, D., Haas, B.J., Tickle, T.L., Trombetta, J.J., Baral, P., Klose, C.S.N., Mahlakõiv, T., Artis, D., Rozenblatt-Rosen, O., Chiu, I.M., Levy, B.D., Kowalczyk, M.S., Regev, A., Kuchroo, V.K., 2017. The neuropeptide NMU amplifies ILC2-driven allergic lung inflammation. Nature 549, 351–356. https://doi.org/10.1038/nature24029

41. Weizman, O.-E., Adams, N.M., Schuster, I.S., Krishna, C., Pritykin, Y., Lau, C., Degli-Esposti, M.A., Leslie, C.S., Sun, J.C., O’Sullivan, T.E., 2017. ILC1 Confer Early Host Protection at Initial Sites of Viral Infection. Cell 171, 795–808.e12. https://doi.org/10.1016/j.cell.2017.09.052

42. Zeis, P., Lian, M., Fan, X., Herman, J.S., Hernandez, D.C., Gentek, R., Elias, S., Symowski, C., Knöpper, K., Peltokangas, N., Friedrich, C., Doucet-Ladeveze, R., Kabat, A.M., Locksley, R.M., Voehringer, D., Bajenoff, M., Rudensky, A.Y., Romagnani, C., Grün, D., Gasteiger, G., 2020. In Situ Maturation and Tissue Adaptation of Type 2 Innate Lymphoid Cell Progenitors. Immunity 53, 775–792.e9. https://doi.org/10.1016/j.immuni.2020.09.002

